# Natural volatiles causing inhibition of mosquito biting behaviors: development of a virtual screening platform predicting antagonists of ORco function for accelerated discovery

**DOI:** 10.1101/2024.06.24.600390

**Authors:** Georgia Kythreoti, Trias Thireou, Christos Karoussiotis, Zafiroula Georgoussi, Panagiota GV Liggri, Dimitrios P Papachristos, Antonios Michaelakis, Vasileios Karras, Spyros E Zographos, Stefan Schulz, Kostas Iatrou

**Author notes:** Department of Science and Mathematics, Deree – The American College of Greece, Athens, Greece (Georgia Kythreoti). Communicating authors: &. Authors with equal contribution (Georgia Kythreoti, Trias Thireou).

## Abstract

Insect olfactory receptors are heteromeric ligand-gated cation channels composed of an obligatory receptor subunit, ORco, and one of many variable subunits, ORx, in as yet undefined molar ratios. When expressed alone *ex vivo*, ORco forms homotetrameric channels gated by ORco-specific ligands acting as channel agonists. Using an insect cell-based system as a functional platform for expressing mosquito odorant receptors *ex vivo,* we identified small molecules of natural origin acting as specific ORco channel antagonists, orthosteric or allosteric relative to a postulated ORco agonist binding site, and causing severe inhibition of olfactory function in mosquitoes. In the present communication, we are reporting on the compilation of common structural features of such orthosteric antagonists and development of a ligand-based pharmacophore whose properties are deemed necessary for binding to the agonist binding site and inhibition of ORco’s biological function. *In silico* screening of an available collection of natural volatile compounds with this pharmacophore resulted in identification of several ORco antagonist hits. Cell-based functional screening of the same compound collection resulted in the identification of several compounds acting as orthosteric and allosteric antagonists of ORco channel function *ex vivo* and inducing anosmic behaviors to *Aedes albopictus* mosquitoes *in vivo.* Comparison of the *in silico* screening results with those of the functional assays revealed that the pharmacophore predicted correctly 7 out of the 8 confirmed orthosteric antagonists and none of the allosteric ones. Because the pharmacophore screen also produced additional hits that did not cause inhibition of the ORco channel function, we generated a Support Vector Machine (SVM) model based on two descriptors of all pharmacophore hits. Training of the SVM on the *ex vivo* validated compound collection resulted in the selection of the confirmed orthosteric antagonists with a very low cross-validation out-of-sample misclassification rate. Employment of the combined pharmacophore-SVM platform for *in silico* screening of a larger collection of olfaction-relevant volatiles produced several new hits. Functional validation of randomly selected hits and rejected compounds from this screen confirmed the power of this virtual screening platform as convenient tool for identifying novel vector control agents. To the best of our knowledge, this study is the first one that combines a pharmacophore with a SVM model for identification of AgamORco antagonists and specifically orthosteric ones.

## INTRODUCTION

Many insect species have the potential to transmit a wide range of pathogens to humans and animals, causing a variety of vector-borne diseases (VBDs). According to the World Health Organization (WHO), VBDs account for more than 17% of all infectious diseases, causing more than 700,000 deaths annually [1]. Therefore, they pose a significant threat to global public and animal health and have substantial socioeconomic impacts. Although effective control of insect disease vectors (IDVs) is crucial, it is also quite challenging. One powerful, effective and safe control method involves the use of long lasting and environmentally friendly repellents and anosmia-inducing agents. These agents interfere with the olfactory capacity of blood-feeding insects thus reducing the possibility of their biting host organisms and transmitting pathogens to them.

Insects rely on their olfactory system to sense volatile chemicals that regulate various behaviors, including social interactions, mate and oviposition site selection, food source location and enemy recognition [2, 3]. Insect odor receptors, expressed in olfactory sensory neurons, are heteromeric ligand-gated cation channels. They are composed of one of many variable subunits, ORx, which confer specificity in the recognition of the odor molecules, and an obligatory receptor subunit, ORco, which is necessary for channel formation and signal transduction [4, 5] in as yet undetermined molar ratios. In contrast to the variable ORx subunits, ORco is highly conserved amongst different insect orders, spanning many hundreds of millions of years of evolution [2, 6, 7]. Moreover, we and others have shown that in the absence of a co-expressed ORx subunit, ORco can form in vitro homotetrameric cation channels [8, 9] whose function may be activated or suppressed by synthetic ORco agonists and antagonists [10–15]. Additionally, ORco antagonists have broad inhibitory activities on the majority of ORs of a variety of insects, and their binding site(s) on ORco may thus serve as “universal” modulatory site(s) for volatile compounds. Given such considerations, we set out to identify new ORco antagonists interrupting insect – host recognition and thus reducing and preventing the spread of VBDs.

Discovery of bioactive molecules through experimental screening of large compound collections is an expensive and time-consuming process. The complexity of this process may be greatly reduced by the availability of appropriate *in vitro* or *ex vivo* functional assays and, even more so, the undertaking of initial virtual screening (VS) steps that use the physicochemical and structural properties of compounds and/or target proteins, along with experimentally verified bio-interaction information, to generate predictive models for identification of candidate bioactive molecules. Hence, VS methods narrow the search space and reduce the time and cost required for a screening project.

Several techniques are currently used for VS, among which the pharmacophore method and machine learning. The pharmacophore is an ensemble of steric and electronic features that ensure optimal supramolecular interactions with a specific biological target structure that may lead to activation or blocking of its biological response [16]. The simplicity and abstract nature of the pharmacophore concept enables the complexity of interactions between ligands and receptors to be reduced to a small set of features [17]. Thus, pharmacophore-based techniques have become an integral part of computer-aided drug design and have been successfully applied for virtual screening, *de novo* design, and lead optimization [18]. Pharmacophore models can be derived from experimentally determined protein-ligand complexes (receptor-based pharmacophores), or from known active compounds (ligand-based pharmacophores).

Machine learning, on the other hand, has established itself as a VS methodology in its own right and is constantly growing in popularity. Both conventional machine learning methods, such as Support Vector Machines (SVMs), and deep learning methods are used [19–21]. A support vector machine (SVM) is a supervised learning algorithm with a growing number of applications in precision medicine and drug discovery [22, 23]. In a SVM binary classification problem, a high dimension decision surface is constructed [24, 25]. Several different kernels are introduced to map the data to the featured space, making SVMs able to handle various nonlinear problems with improved generalization characteristics.

In the present study, we are reporting on the development of a two-step VS protocol that achieves the goal of accelerating the discovery process for new bioactive molecules that prevent mosquitoes from obtaining blood meals from their hosts by virtue of acting as antagonists of the ORco channel in mosquitoes. In the first step, a pharmacophore model was constructed based on a set of small ligands, that we have previously determined to function as specific ORco channel antagonists, orthosteric or allosteric relative to the ORcoRAM2 agonist binding site [14, 26], causing severe inhibition of olfactory function in mosquitoes [26, 27]. Sequentially, a SVM model was applied to refine the results and to better prioritize the compounds for experimental validation. The usefulness of the specific VS protocol is assessed by *ex vivo* assays using a previously developed cell-based functional platform [10–15].

## RESULTS

### Structural features of orthosteric ORco antagonists and pharmacophore design

Our previous studies on a limited collection comprising 54 volatile organic compounds (VOCs) of natural origin have led to the identification of several ORco ligands, which acted as antagonists of the homomerized ORco subunit [14, 26, 28]. Some of the identified antagonists were also shown to possess powerful repellent activities for different mosquito species [26, 27]. Moreover, based on competition assays against OrcoRAM2, a previously characterized ORco agonist predicted to bind to each ORco subunit of a homotetrameric ORco channel at a hypothesized site [8, 9, 29, 30], the identified antagonists were classified as orthosteric or allosteric relative to the OrcoRAM2 binding site. The previously identified orthosteric and allosteric Orco ligands and their structural formulae and chemical classes are shown in **Table 1**.

**Table 1.**
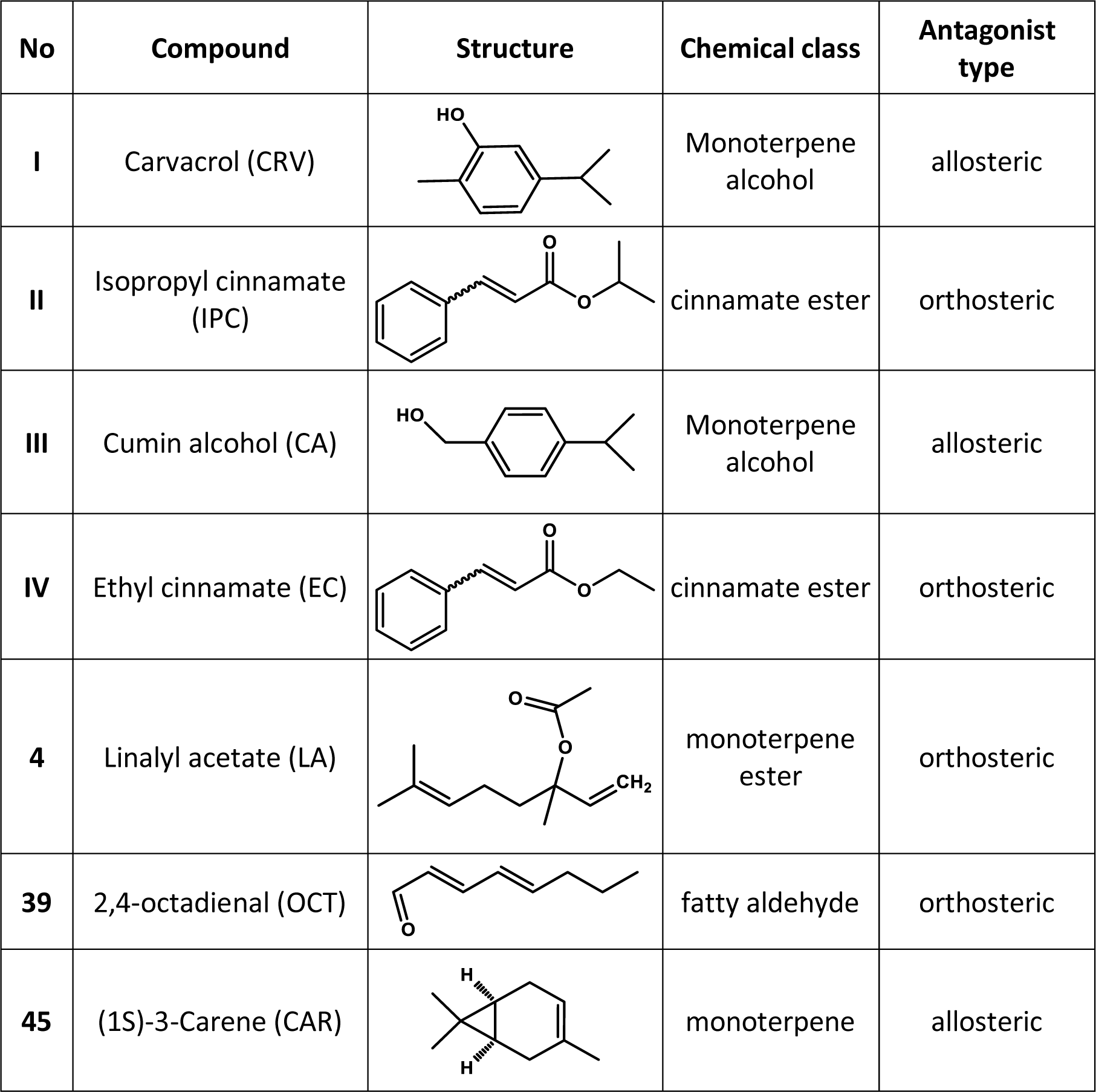
Previously identified ORco orthosteric and allosteric antagonists. Structural features of previously identified AgamORco orthosteric and allosteric antagonists. Compound numbers are the same as those presented in [26].

| No | Compound | Structure | Chemical class | Antagonist type |
| --- | --- | --- | --- | --- |
| I | Carvacrol (CRV) |  | Monoterpene alcohol | allosteric |
| II | Isopropyl cinnamate (IPC) |  | cinnamate ester | orthosteric |
| III | Cumin alcohol (CA) |  | Monoterpene alcohol | allosteric |
| IV | Ethyl cinnamate (EC) |  | cinnamate ester | orthosteric |
| 4 | Linalyl acetate (LA) |  | monoterpene ester | orthosteric |
| 39 | 2,4-octadienal (OCT) |  | fatty aldehyde | orthosteric |
| 45 | (1S)-3-Carene (CAR) |  | monoterpene | allosteric |

In an effort to accelerate the further discovery of compounds capable of antagonizing ORco function, in terms of identification of orthosteric ORco antagonist candidates (hits) in available VOC collections of natural or synthetic origin, we sought to develop a ligand-based pharmacophore that could describe orthosteric antagonist features necessary for blocking ORco’s biological response. If successful, the specific pharmacophore could be employed as probe for an initial virtual screening of available compound collections prior to carrying out relevant functional screens.

Using as a training set the previously characterized collection of 54 VOCs, which included 4 positive examples (the orthosteric antagonists, shown in **Table 1)** and 50 negative ones (the 3 allosteric antagonists shown in **Table 1** and 47 inactive compounds shown in **Table S1**), a ligand-based pharmacophore has been developed that described the 3D arrangement of orthosteric antagonist features necessary for blocking ORco’s biological response. The specific pharmacophore model has been required to match all orthosteric input molecules, while keeping the number of false positives (allosteric antagonists and inactive compounds) at a minimum. Four features (**Figure 1**) were found to meet these requirements best. These included one atom-centered hydrophobic feature “HydA”, two centroid hydrophobic features “Hyd” and one projected location of potential H-bond donors “Acc2”. Hydrogen bond Acc2 projected annotations are added to those heavy atoms that qualify as H-bond acceptors and are given Acc annotations (MOE 2016, Pharmacophore Annotation Schemes; see Materials & Methods). The statistical significance of our model was estimated at -4.4626 (MOE 2016, The Pharmacophore Elucidator; see Materials & Methods).

**Figure 1.**
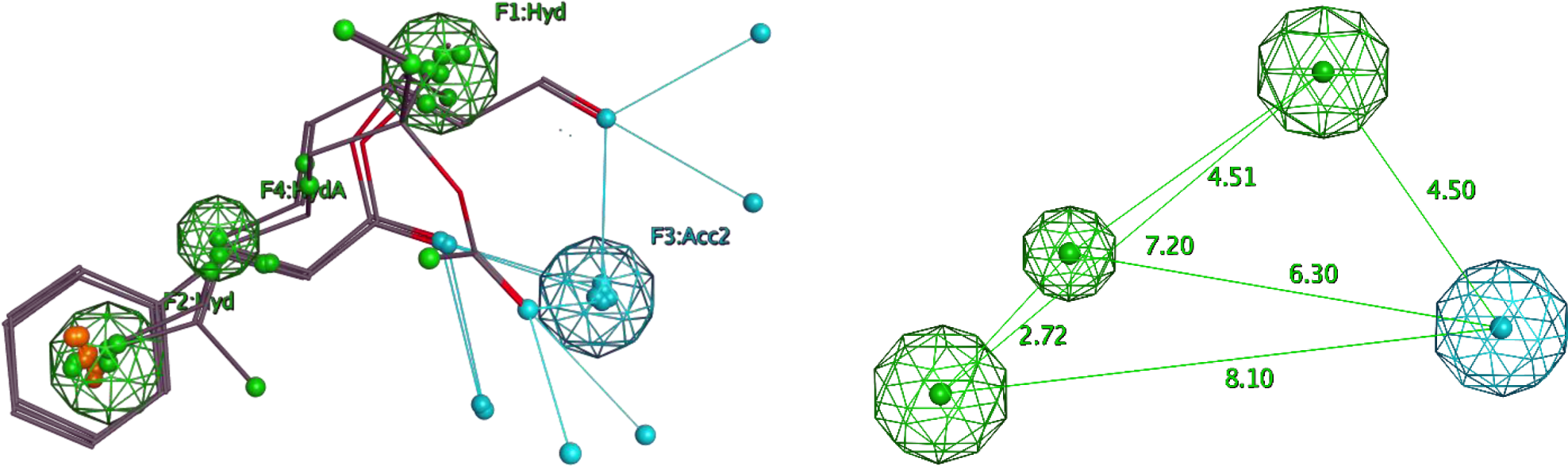
The pharmacophore model. *<u>Left</u>*: The ligand-based pharmacophore and the 4 orthosteric antagonists of **Table 1** (Isopropyl cinnamate, Ethyl cinnamate, Linalyl acetate, 2,4-octadienal) used to generate it. The features include one atom-centered hydrophobic feature HydA (green), two centroid hydrophobic features Hyd (green) and one projected location of potential H-bond donors Acc2 (blue). *<u>Right:</u>* Spacing distances between the specific pharmacophore features.

The results of the training process of the specific pharmacophore model on the collection of 54 VOCs of **Table S1**, are shown in **Table 2**.

**Table 2.**
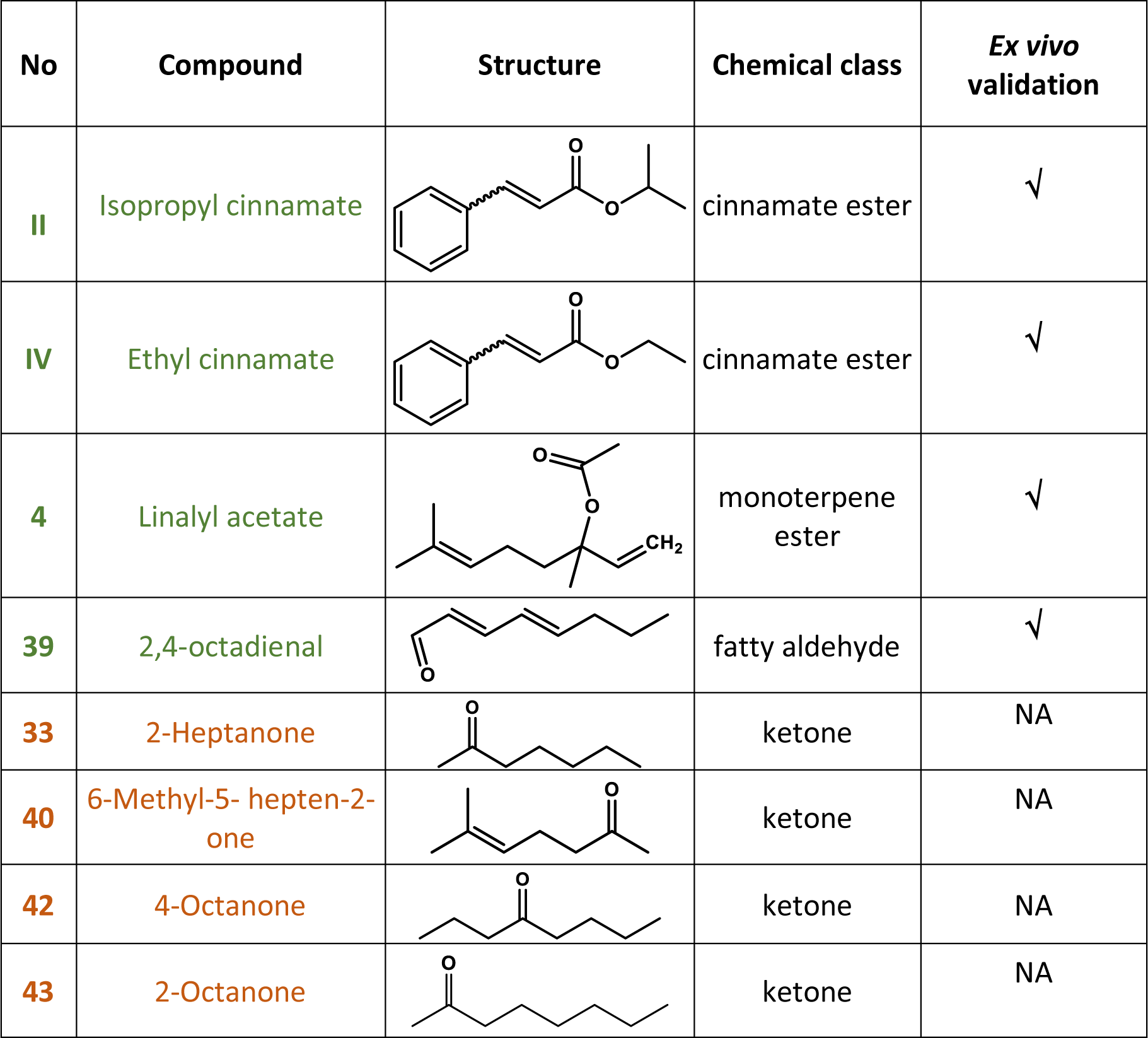
ORco orthosteric antagonist hits and *ex vivo* validation. The training set for the selected pharmacophore model consisted of the four confirmed orthosteric antagonists shown in Table 1 and fifty negative examples (3 allosteric and 47 inactive) shown in Table 1 and **Table S1**. Compound numbering is as per [26] and **Table S1**; functionally validated (bioactive) hits are shown in green, while inactive ones are shown in orange; **√**: >40% inhibition; NA: not active (<40% inhibition).

| No | Compound | Structure | Chemical class | *Ex vivo* validation |
| --- | --- | --- | --- | --- |
| II | Isopropyl cinnamate |  | cinnamate ester | ✓ |
| IV | Ethyl cinnamate |  | cinnamate ester | ✓ |
| 4 | Linalyl acetate |  | monoterpene ester | ✓ |
| 39 | 2,4-octadienal |  | fatty aldehyde | ✓ |
| 33 | 2-Heptanone |  | ketone | NA |
| 40 | 6-Methyl-5-hepten-2-one |  | ketone | NA |
| 42 | 4-Octanone |  | ketone | NA |
| 43 | 2-Octanone |  | ketone | NA |

As may be seen in **Table 2**, screening of the training set with the selected pharmacophore resulted in the expected recognition of the four previously identified ORco orthosteric antagonists, compounds II, IV, 4 and 39 (ORco *ex vivo* inhibition of >40%; [26]. In addition, however, the screening identified four more hits (compounds 33, 40, 42, 43), which either did not display any antagonist activities in our *ex vivo* activity screens (compound 42) or caused only minor inhibition of ORco activity, in the order of 15-20% (compounds 33, 40 and 43) [26]. The remaining 46 compounds, including the three previously identified ORco allosteric antagonists shown in **Table 1**, were not selected by the pharmacophore.

### *In silico* screening of a new VOC collection and *ex vivo* validation

To examine the performance of the specific pharmacophore model as a probe for a virtual screen of a previously “unseen” set of data, we screened *in silico* an additional collection of 49 natural VOCs (**Table S2**). The pharmacophore model yielded a total of 24 hits, shown in **Table 3**.

**Table 3.**
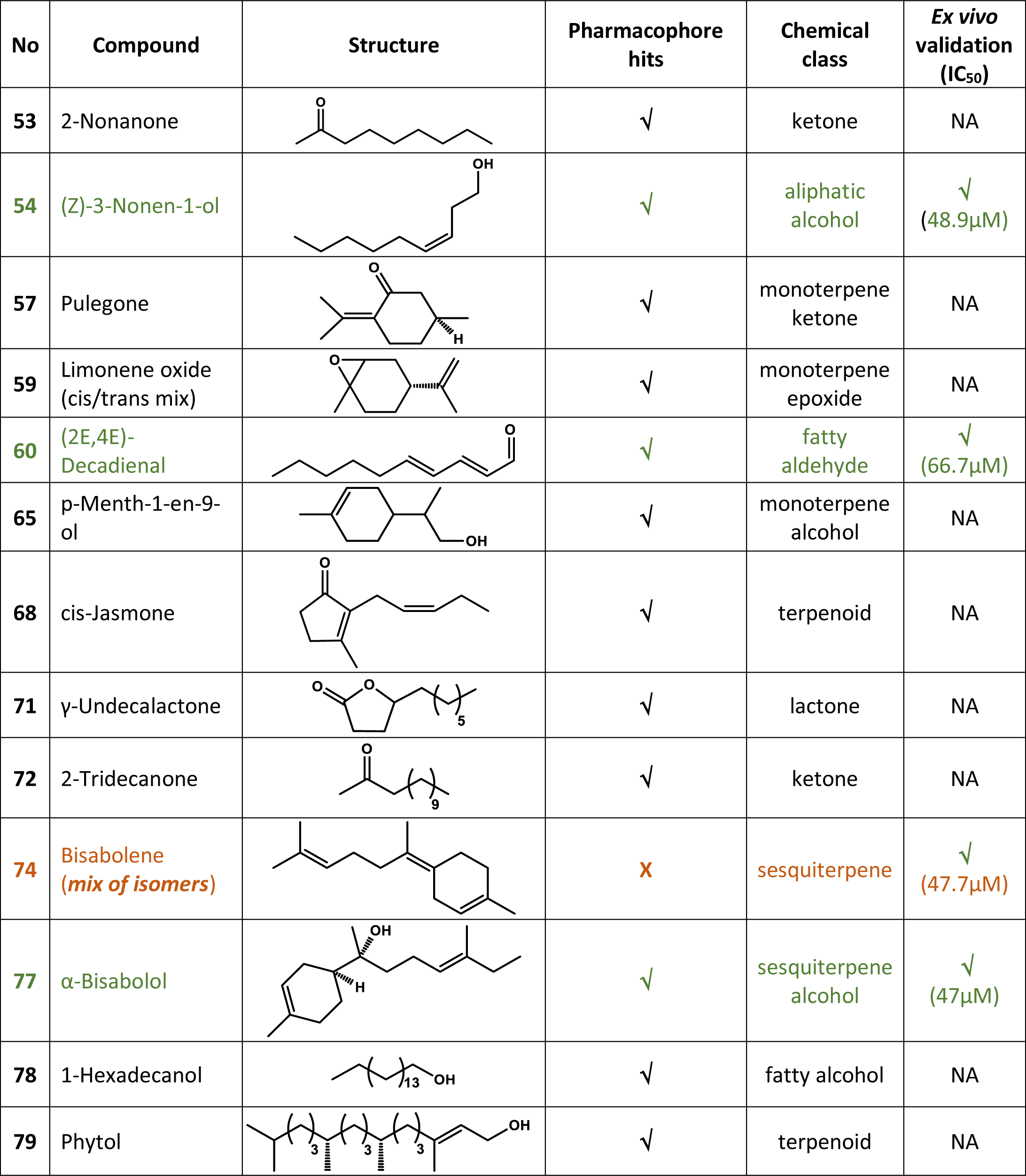
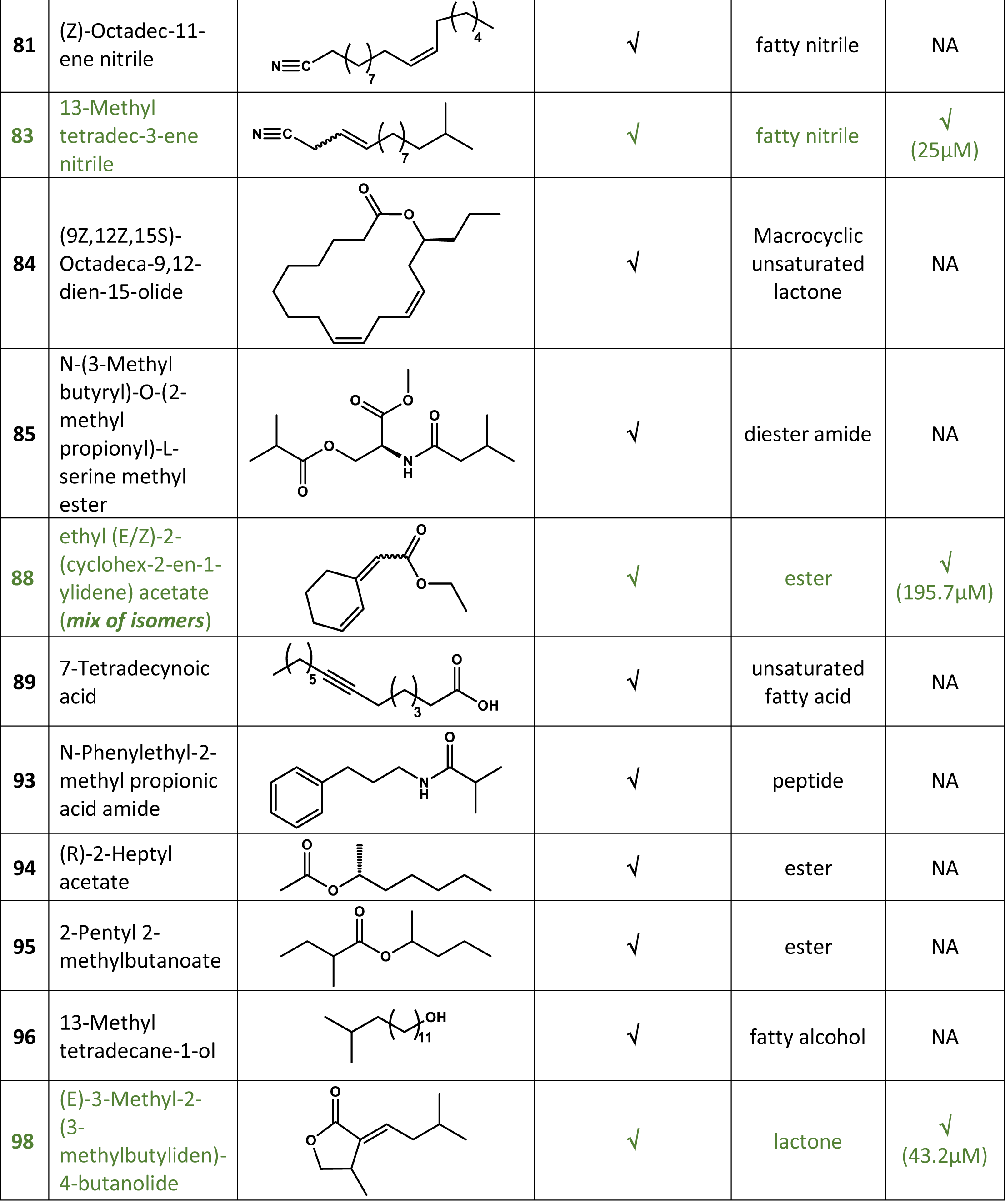
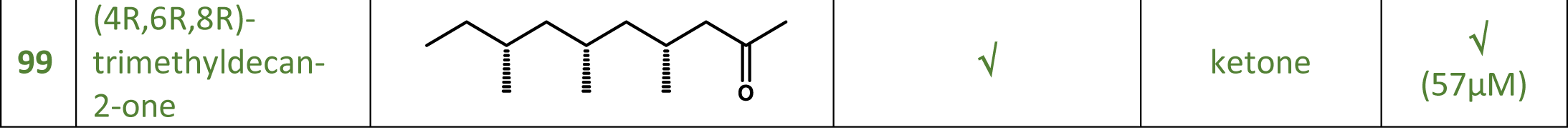
*In silico pharmacophore screening* and *ex vivo* functional validation of a new compound collection. Antagonists have been defined as VOCs causing at least 40% inhibition of ORco activity in the *ex vivo* assay as per tests shown in **Figure 2**. X: not detected by the *in silico* screen; **√**: pharmacophore antagonist hits; NA: Not active or less than 40% maximum inhibition in the competition assay. IC_50_: Concentration of 50% inhibition in the presence of 100μM ORco agonist. Green color indicates *ex vivo* active pharmacophore hits whereas orange color indicates the *ex vivo* active compound that escaped detection by the pharmacophore.

| No | Compound | Structure | Pharmacophore hits | Chemical class | *Ex vivo* validation (IC<sub>50</sub>) |
| --- | --- | --- | --- | --- | --- |
| 53 | 2-Nonanone |  | ✓ | ketone | NA |
| 54 | (Z)-3-Nonen-1-ol |  | ✓ | aliphatic alcohol | ✓<br>(48.9µM) |
| 57 | Pulegone |  | ✓ | monoterpene ketone | NA |
| 59 | Limonene oxide (cis/trans mix) |  | ✓ | monoterpene epoxide | NA |
| 60 | (2E,4E)-Decadienal |  | ✓ | fatty aldehyde | ✓<br>(66.7µM) |
| 65 | p-Menth-1-en-9-ol |  | ✓ | monoterpene alcohol | NA |
| 68 | cis-Jasmone |  | ✓ | terpenoid | NA |
| 71 | γ-Undecalactone |  | ✓ | lactone | NA |
| 72 | 2-Tridecanone |  | ✓ | ketone | NA |
| 74 | Bisabolene (mix of isomers) |  | X | sesquiterpene | ✓<br>(47.7µM) |
| 77 | α-Bisabolol |  | ✓ | sesquiterpene alcohol | ✓<br>(47µM) |
| 78 | 1-Hexadecanol |  | ✓ | fatty alcohol | NA |
| 79 | Phytol |  | ✓ | terpenoid | NA |

| No | Compound | Structure | Pharmacophore hits | Chemical class | Ex vivo validation (IC<sub>50</sub>) |
| --- | --- | --- | --- | --- | --- |
| 81 | (Z)-Octadec-11-ene nitrile |  | √ | fatty nitrile | NA |
| 83 | 13-Methyl tetradec-3-ene nitrile |  | √ | fatty nitrile | √<br>(25μM) |
| 84 | (9Z,12Z,15S)-Octadeca-9,12-dien-15-olide |  | √ | Macrocyclic unsaturated lactone | NA |
| 85 | N-(3-Methyl butyryl)-O-(2-methyl propionyl)-L-serine methyl ester |  | √ | diester amide | NA |
| 88 | ethyl (E/Z)-2-(cyclohex-2-en-1-ylidene) acetate (mix of isomers) |  | √ | ester | √<br>(195.7μM) |
| 89 | 7-Tetradecynoic acid |  | √ | unsaturated fatty acid | NA |
| 93 | N-Phenylethyl-2-methyl propionic acid amide |  | √ | peptide | NA |
| 94 | (R)-2-Heptyl acetate |  | √ | ester | NA |
| 95 | 2-Pentyl 2-methylbutanoate |  | √ | ester | NA |
| 96 | 13-Methyl tetradecane-1-ol |  | √ | fatty alcohol | NA |
| 98 | (E)-3-Methyl-2-(3-methylbutyliden)-4-butanolide |  | √ | lactone | √<br>(43.2μM) |
| 99 | (4R,6R,8R)-trimethyldecan-2-one |  | ✓ | ketone | ✓<br>(57μM) |

**Figure 2.**
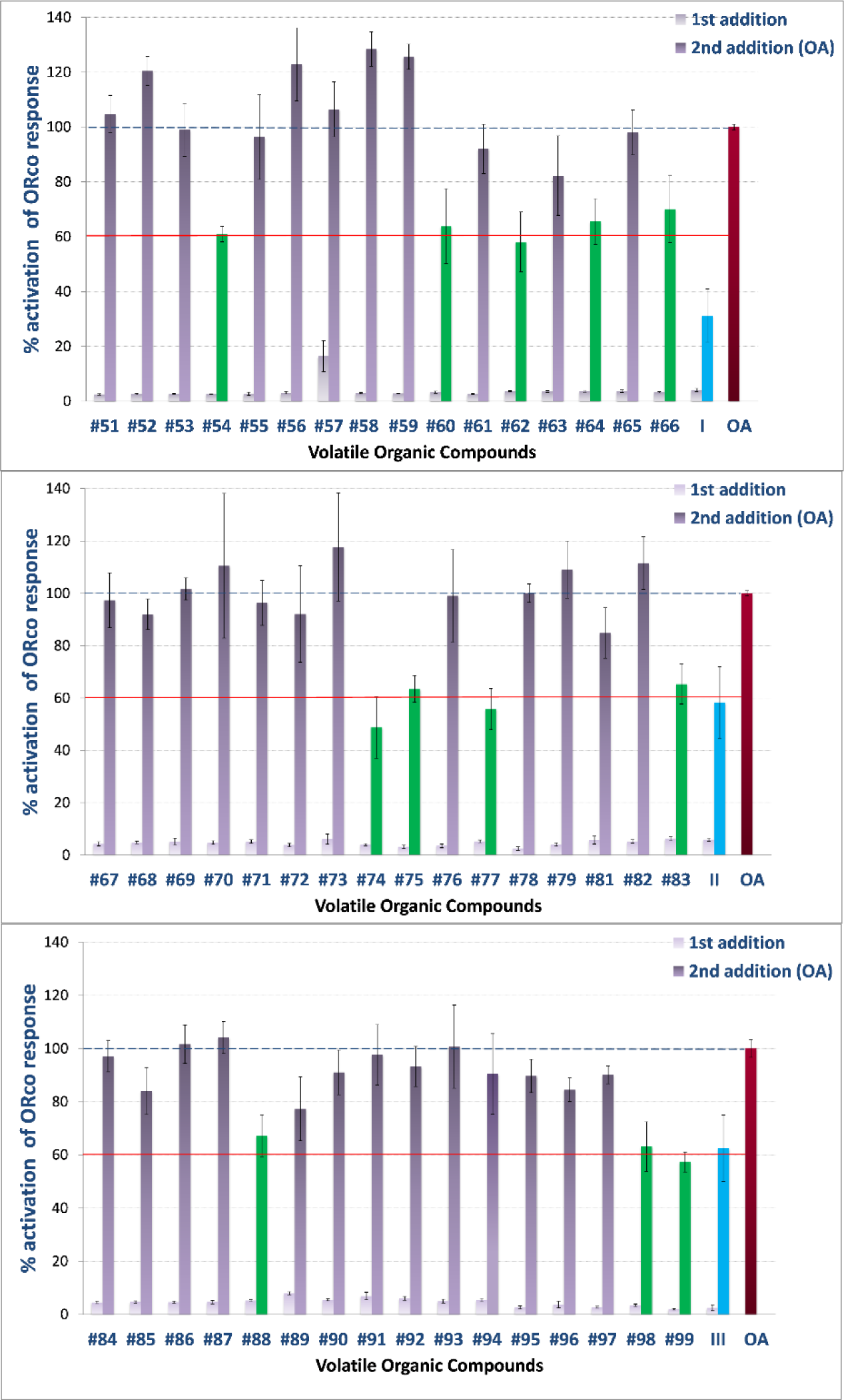
*Ex vivo* screening results. All compounds were tested at a final concentration of 100μM. The primary compound additions (white bars) did not induce significant ORco channel function, while secondary additions of the OA (ORcoRAM2) to wells containing primary additions of functionally inactive compounds produced responses (grey bars) equal to at least 80% of the full response obtained in the control wells (OA only added, set as 100%; red bar at right of each panel). ORco antagonist hits (green bars) produced significantly lower secondary responses, set arbitrarily at ⩽60% of the normal channel response, upon OA addition. Arabic numbers correspond to those of the compounds listed in Table S2, while roman numbers are those of the previously characterized ORco antagonists (blue bars) shown in **Table 1**. Error bars indicate mean±SE.

Concurrently, this collection was functionally screened *ex vivo* using the previously described cell-based platform for determining the % inhibition in ORco agonist activity. This screen uncovered the presence of 12 active compounds in this collection of natural VOCs, which caused a substantial, equal or greater than 40%, degree of inhibition to the activity of the homomeric ORco channels. The results of the cell-based activity screen are shown in **Figure 2**. The structure of the 7 *ex vivo* active pharmacophore hits is shown superimposed onto the pharmacophre model in **Supplementary Figure S1**.

The bioactive VOCs were also subjected to competition tests against the ORco agonist OrcoRAM2 to deduce IC_50_ values and distinguish orthosteric from allosteric antagonists. The competition assays were carried out using as competitors three different concentrations the ORco agonist OrcoRAM2 (50, 100 and 150 μM). These assays, representatives of which are shown in Figure 3, provided the measure of inhibitory activities, IC_50_ values, for the confirmed antagonists, vis-à-vis the *ex vivo* ORco activity normally induced by the presence of 100 μM of OA. Secondly, they allowed the distinction between ORco allosteric and orthosteric antagonists relative to the ORco agonist binding site.

**Figure 3.**
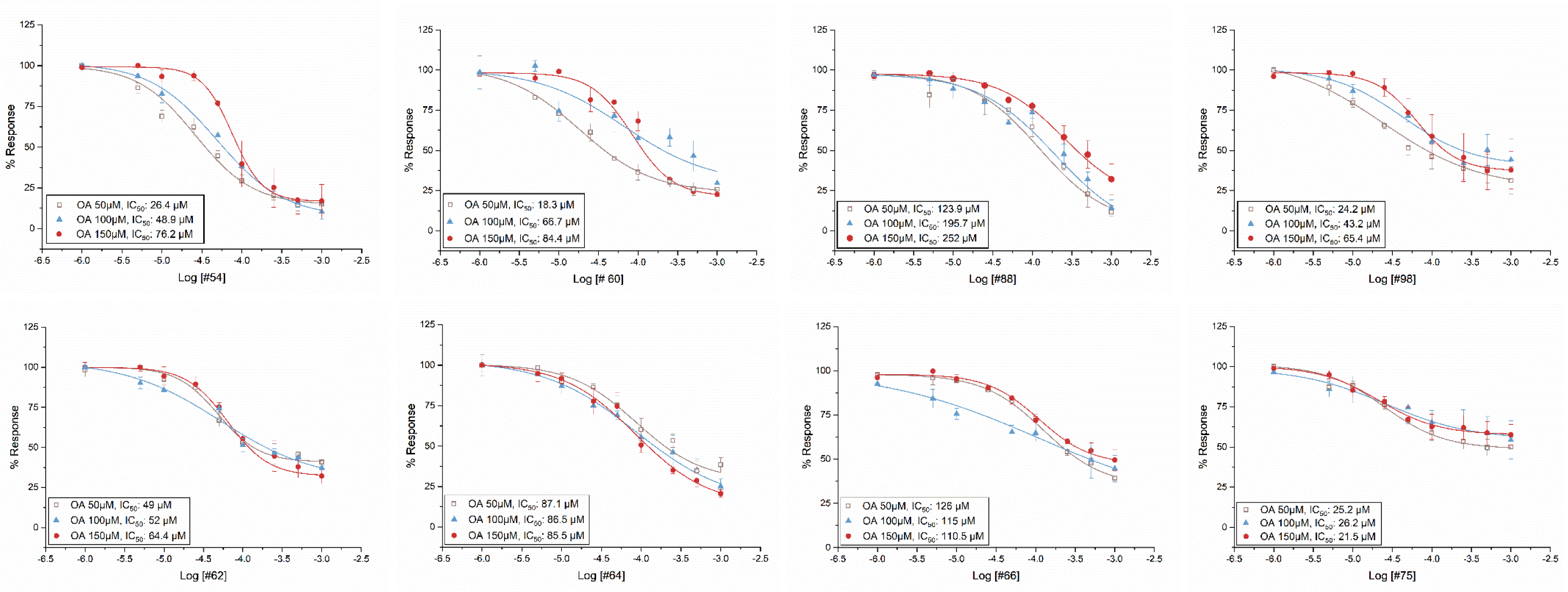
Competition plots for 8 active compounds [orthosteric (upper panels) and allosteric antagonists (lower panels)] showing the % Response as a function of ligands concentration in the presence of 50, 100 and 150μΜ of ORcoRAM2. (For additional data on pIC50 and R^2^, please see **Table S3**)

All *ex vivo* validated orthosteric and allosteric antagonists, 8 and 4, respectively, present in the new, virtually screened VOC collection, 8 and 4, respectively, together with their IC_50_ values, are listed in **Tables 3** (green colored compounds) and **4**.

The 24 pharmacophore hits (**Table 3**) included all but one (#74) of the eight orthosteric antagonists identified through the cell-based activity screening and competition assays presented in Figures 2 and **3** (compounds #54, 60, 77, 83, 88, 98 and 99). None of the allosteric antagonists shown in **Table 4** were identified as antagonist hits by the pharmacophore. Thus, the sensitivity of the pharmacophore model for *in silico* prediction of actual orthosteric antagonists present in the specific collection of natural VOCs (**Table S3**) has been an impressive 0.88. However, the remaining 17 hits shown in **Table 3** were found to be either not active against ORco in the *ex vivo* assays or to inhibit Orco activity by substantially less than the previously defined useful inhibition cutoff point of 40% (Figure 2). Accordingly, the specificity of the pharmacophore screen has been 0.59 (see Materials & Methods), a value that may be unsustainable in terms of experimental effort, especially for virtual screening of large libraries. Overall, the performance of the pharmacophore model described above was evaluated using the Power metric (PM) value [31], since this value might estimate better the performance of a virtual screening when few experiments can be carried out. The PM value for the pharmacophore VS was equal to 0.68, leaving room for improvement. For this reason, a second filtering step was added to the *in silico* screening pipeline.

**Table 4.**
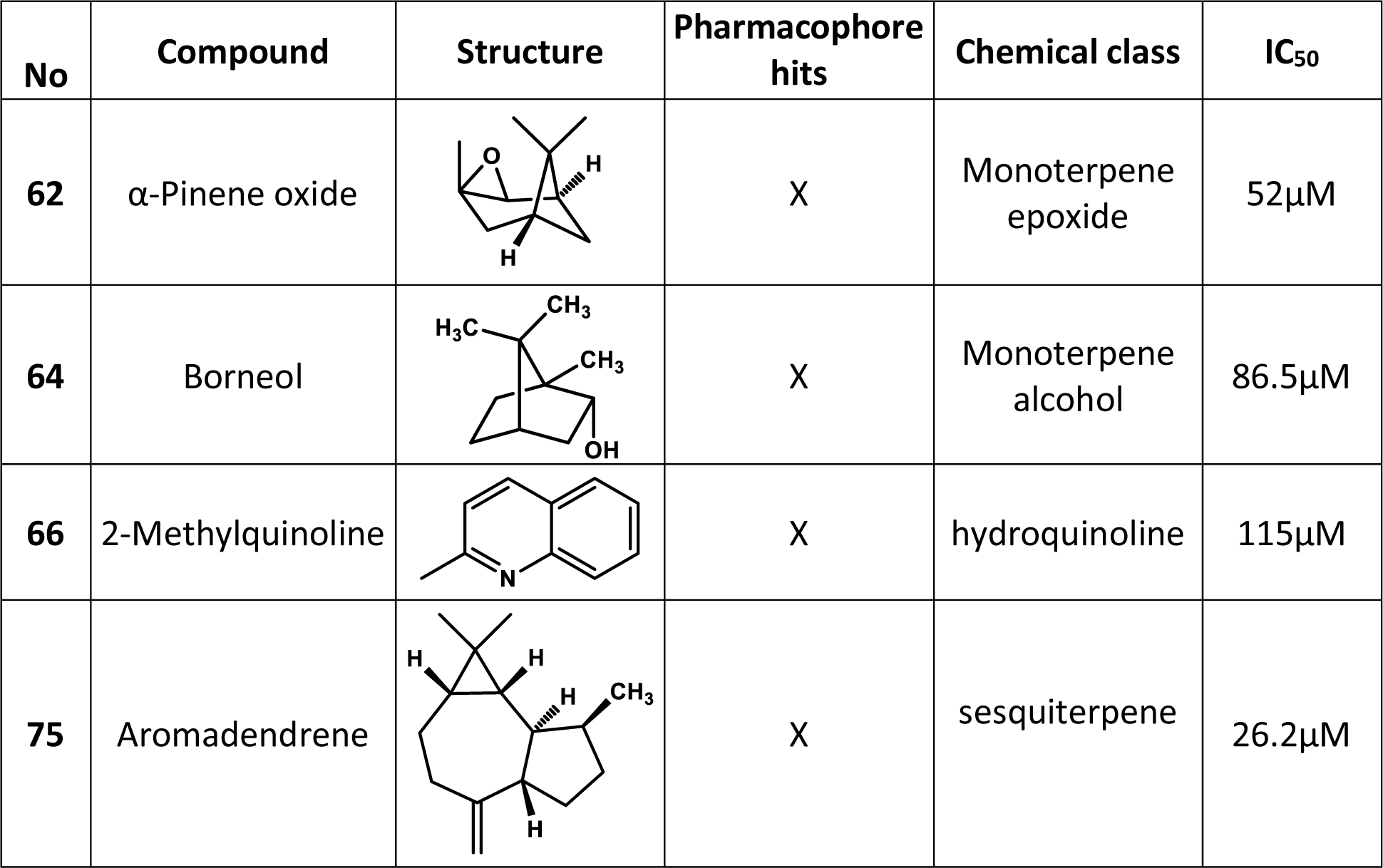
*Ex vivo* validated ORco allosteric antagonists. Structural features and chemical classes of identified AgamORco allosteric antagonists. IC_50_ values presented, in the presence of 100 μM ORco agonist.

| No | Compound | Structure | Pharmacophore hits | Chemical class | IC<sub>50</sub> |
| --- | --- | --- | --- | --- | --- |
| 62 | α-Pinene oxide | | X | Monoterpene epoxide | 52μM |
| 64 | Borneol | | X | Monoterpene alcohol | 86.5μM |
| 66 | 2-Methylquinoline | | X | hydroquinoline | 115μM |
| 75 | Aromadendrene | | X | sesquiterpene | 26.2μM |

### Generation of a SVM model based on 2D descriptors for improved predictions

To improve on the reliability of predictions for the identification of orthosteric antagonist hits, we trained several SVM models using the 32 pharmacophore hits shown in **Tables 2** (8 hits) and **3** (24 hits). The set of 2D descriptors has been calculated in MOE (see Materials and Methods). The descriptor pairs that resulted in the best SVM model included the KierA2 and SlogP_VSA1 structural features. KierA2 or second alpha modified shape index is a topological descriptor that encodes the branching of a molecule. In general, for straight chain molecules, KierA2=A-1 (where A is the atom count). SlogP_VSA1, on the other hand, describes the sum of the accessible van der Waals surface area for each atom whose logarithm of the octanol/water partition coefficient is in the range (-0.4 to -0.2] or, in other words, the extent of hydrophobic or hydrophilic effects on the surface area of the molecule. The SVM model with the lowest out-of-sample misclassification rate was selected and optimized, yielding a cross-validation loss equal to 0.032. The results of applying the selected SVM filters on the 32 pharmacophore hits are detailed in **Table 5** and shown diagrammatically in Figure 4.

**Table 5.** Results of the selected SVM classification of the 32 pharmacophore hits. The table lists the orthosteric antagonist hits identified by the pharmacophore model after the application of the selected SVM model on the training set of compounds. The *ex vivo* active compound #99 was laid outside the decision boundaries of the SVM model (**Figure 4**) and was defined as a false-negative result yielding a cross-validation loss of 0.032. NA: not or marginally active (≤40% inhibition) in the *ex vivo* assays.

| Compound | Cell-based activity (IC<sub>50</sub> in μM) | KierA2 | SlogP_VSA1 | within SVM decision boundaries |
| --- | --- | --- | --- | --- |
| II | 41.7 | 4.5847445 | 7.7454643 | Yes |
| IV | 64.5 | 4.4210858 | 7.7454643 | Yes |
| 4 | 67.7 | 5.2678456 | 7.7454643 | Yes |
| 33 | NA | 4.4425101 | 5.6876111 | No |
| 39 | 59.8 | 5.5963559 | 0 | Yes |
| 40 | NA | 3.809427 | 5.6876111 | No |
| 42 | NA | 5.4008284 | 5.6876111 | No |
| 43 | NA | 5.4008284 | 5.6876111 | No |
| 53 | NA | 6.368185 | 5.6876111 | No |
| 54 | 48.9 | 8.1811314 | 0 | Yes |
| 57 | NA | 2.9886453 | 5.6876111 | No |
| 59 | NA | 2.0609839 | 0 | No |
| 60 | 66.7 | 7.5834055 | 0 | Yes |
| 65 | NA | 3.795996 | 0 | No |
| 68 | NA | 3.8128579 | 5.6876111 | No |
| 71 | NA | 5.9534798 | 7.7454643 | No |
| 72 | NA | 10.287 | 5.6876111 | No |
| 77 | 47 | 5.4362946 | 0 | Yes |
| 78 | NA | 16 | 0 | No |
| 79 | NA | 13.104808 | 0 | No |
| 81 | NA | 16.544603 | 0 | No |
| 83 | 25 | 11.143562 | 0 | Yes |
| 84 | NA | 12.104386 | 7.7454643 | No |
| 85 | NA | 8.1214361 | 20.749712 | No |
| 88 | 195.7 | 5.1365461 | 7.7454643 | Yes |
| 89 | NA | 11.143562 | 7.7454643 | No |
| 93 | NA | 5.4330149 | 5.2587838 | No |
| 94 | NA | 6.0677109 | 7.7454643 | No |
| 95 | NA | 5.9571776 | 7.7454643 | No |
| 96 | NA | 13.066667 | 0 | No |
| 98 | 43.2 | 3.932668 | 7.7454643 | Yes |
| 99 | 57 | 6.7910275 | 5.6876111 | No |

**Figure 4.**
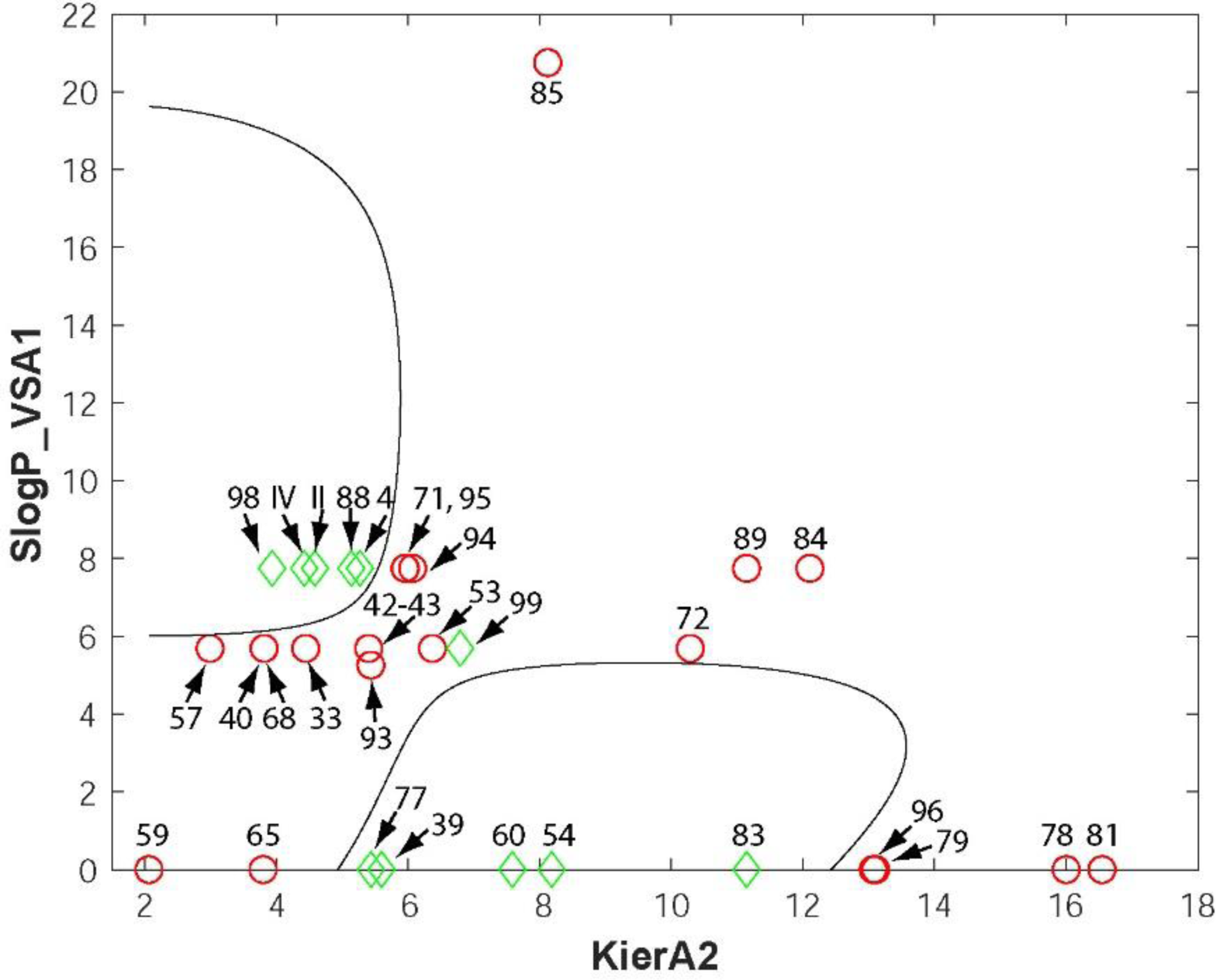
The selected SVM model. The SVM model was trained on all pharmacophore hits (**Tables 2 and3**) using the 2D descriptors KierA2 and SlogP_VSA1 (MOE software). Ex vivo active orthosteric antagonists are represented by green diamonds whereas non-active compounds are indicated by red circles. Compounds with coordinates (KierA2, SlogP_VSA1) that lie within the areas delineated by the decision boundaries (solid lines) are predicted to be orthosteric antagonists of ORco.

The data points of the training classes together with the decision boundaries that separate them in the feature space are visualized in the classification map shown in Figure 4. The RBF kernel handled the non-linearly separable data creating curved decision boundaries.

### VOCs antagonizing ORco function act as spatial, mosquito anosmia-like inducing agents

The functionalities of the new *ex vivo*-validated ORco antagonists, orthosteric and allosteric ones, except those for #54 [(Z)-3-Nonen-1-ol], which was found to have a high IC_50_ value of (178μM; **Table 3**) and #99 due to unavailability of sufficient quantities, were subsequently assessed *in vivo* against *Aedes albopictus* as previously described [26, 32] at different concentrations ranging from a high of 200 to a low of 50 nmole/cm^2^. At such concentrations, all *ex vivo* validated antagonists were found to cause *in vivo* inhibition in the numbers of mosquitoes that landed on the exposed hand areas to various extents (data not shown). Seeking potent repellents, compounds showing significant repellency (RI >50%) at the dose of 50 nmole/cm^2^ were subsequently tested at an even lower dose of 10 nmole/cm^2^. Thus, while compounds #39, #54, #77 and the allosteric antagonist #62 that exhibited mild repellent activity (RI 30%-50%; data not shown) were excluded from further testing, 7 new antagonists displaying high activities in the preliminary *in vivo* tests, 4 orthosteric (#60, 83, 88 and 98) and 3 allosteric ones (#64, 66 and 75) were assessed at the low compound dose of 10 nmole/cm^2^ (Figure 5).

**Figure 5.**
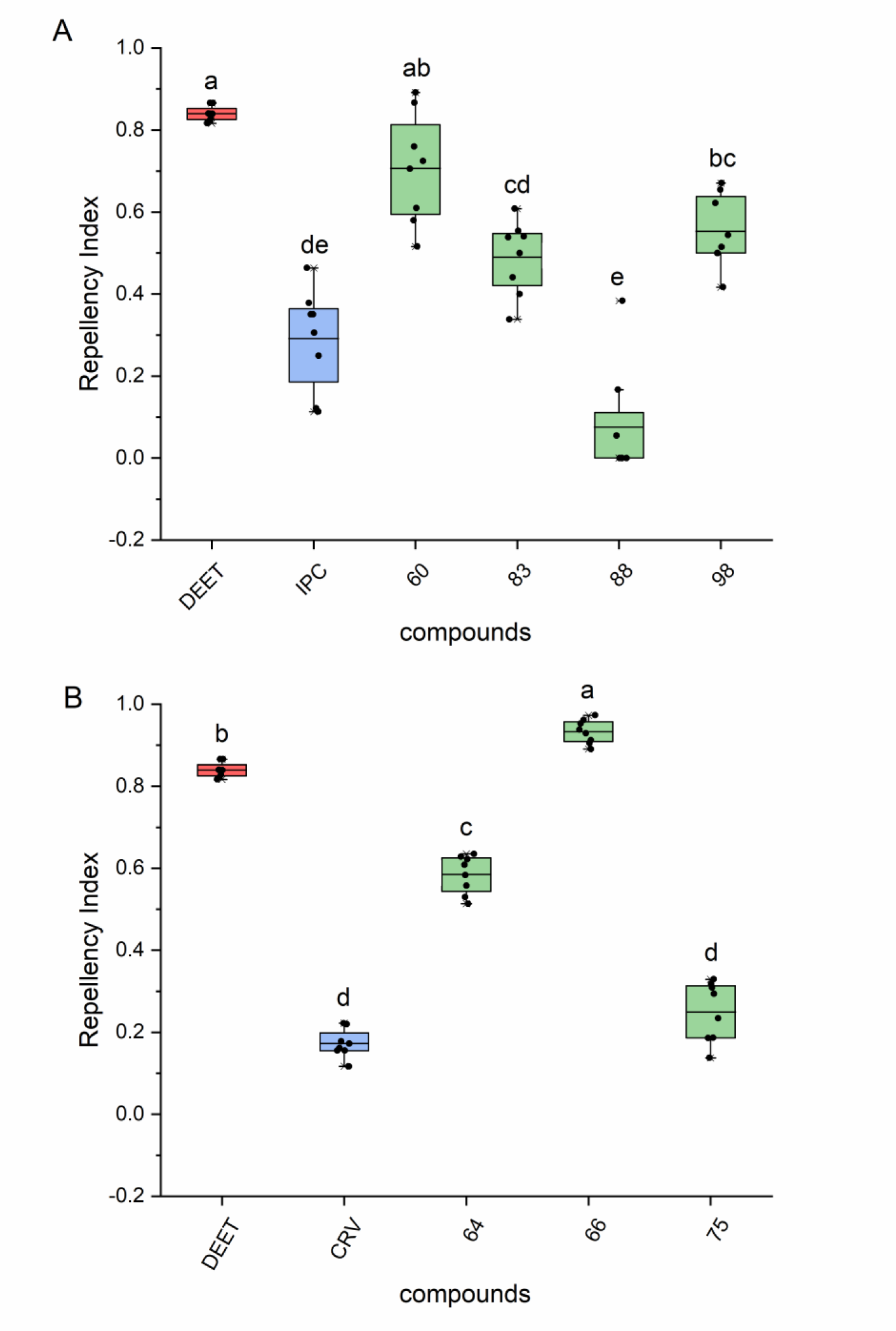
Box plots depicting repellency indices (RIs) against *Aedes albopictus* mosquitoes in “Hand in a cage” repellence assays. **(Α)** Selected orthosteric and **(Β)** allosteric antagonists (green) and the widely used insect repellent DEET (red) were examined using 10nmole of each tested compound per cm^2^ of exposed hand area (240nmole/24cm^2^ total exposed area). The previously characterized antagonists isopropyl cinnamate (IPC; blue) and carvacrol (CRV; blue) [26] served as controls for the tested orthosteric and allosteric antagonists, respectively. The box plots represent the mean values with upper and lower quartiles, and the range of outliers within 1.5IQR are indicated by error bars. Compound identities are listed in **Table S1** and **S2**. Different letters (a, b, etc.) indicate statistically significant differences between tested compounds (p<0.05), Mann–Whitney U test with Bonferroni correction (adjusted p values a=0.005 and a=0.003 for the orthosteric and allosteric group, respectively).

As can be seen from the results presented in Figure 5 **and Table S4**, even at the very low dose of 10 nmole of compound per cm^2^ of naked hand area, mosquitoes exposed to all but one (#88) tested ORco orthosteric antagonists, identified through the combined employment of *in silico* and *ex vivo* screening, were found to display noticeably reduced attraction responses to the human smell emissions. Of particular note has been the orthosteric antagonist #60 [(2E,4E)-Decadienal; RI=0.71± 0.05] and allosteric antagonist #66 (2-Methylquinoline; RI=0.93±0.01), which caused aversion to the hand emissions comparable to that of DEET (RI=0.84±0.01).

### *In silico* screening for additional ORco orthosteric antagonists

To examine the validity of the optimized 2-step *in silico* screening protocol, we virtually screened a new collection of 241 compounds, most of them olfaction-relevant volatiles [**Supplemental spreadsheet**; [33, 34]] for the presence of additional orthosteric antagonists of ORco. Initial application of the specific pharmacophore model on this VOC collection resulted in the identification of 100 hits (Figure 6), while subsequent application of the SVM filter excluded another 56 compounds. Thus, the two-step protocol predicted the presence of 44 putative orthosteric antagonists in this compound collection (Figure 6).

**Figure 6.**
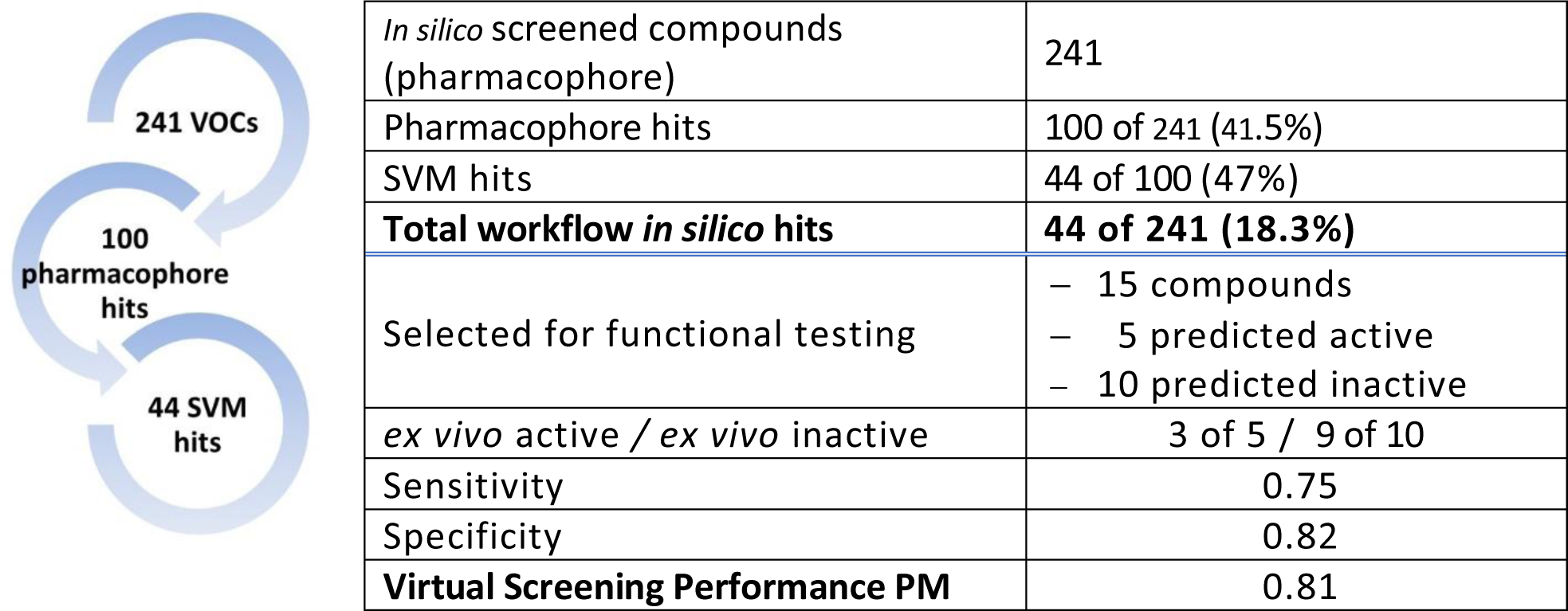
*in silico* screening of a new VOC library [34] for orthosteric Orco antagonists. Starting from 241 VOCs, the pharmacophore identified 100 hits, 44 of which were retained by the SVM filter.

Subsequently, a set of 15 compounds comprised of 5 randomly selected *in silico* hits (putative orthosteric antagonists) and 10 randomly selected workflow-rejected compounds mapping was selected for *ex vivo* functional testing. The mapping of the selected 15 compounds, relative to the established SVM classification map boundaries, is shown diagrammatically in Figure 7.

**Figure 7.**
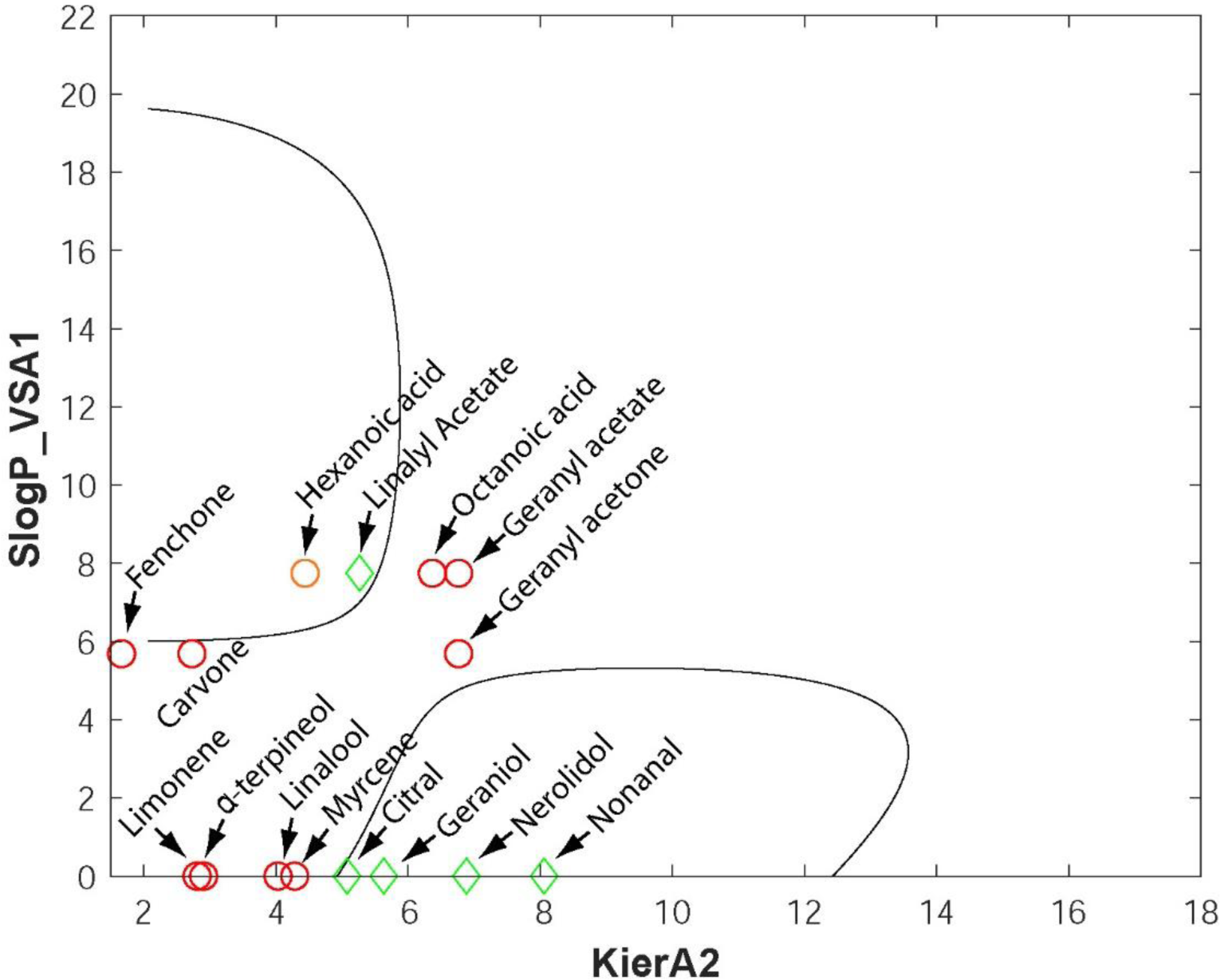
The SVM classification maps. The locations of the five workflow-retained hits (green diamond) and the ten workflow-rejected compounds (red circles) (see also Table 6) are shown in the diagram in the context of their inclusion within or exclusion from the defined SVM boundaries. Hexanoic acid, which was rejected by the pharmacophore model, is indicated by an orange circle inside the upper SVM boundary of bioactive hits. The *ex vivo* activities of the fifteen compounds are shown in **Figure 8**.

The results of the *ex vivo* functional testing for the 15 selected representatives, whose SVM mapping coordinates have been presented in Figure 7 above, are shown in Figure 8.

**Figure 8.**
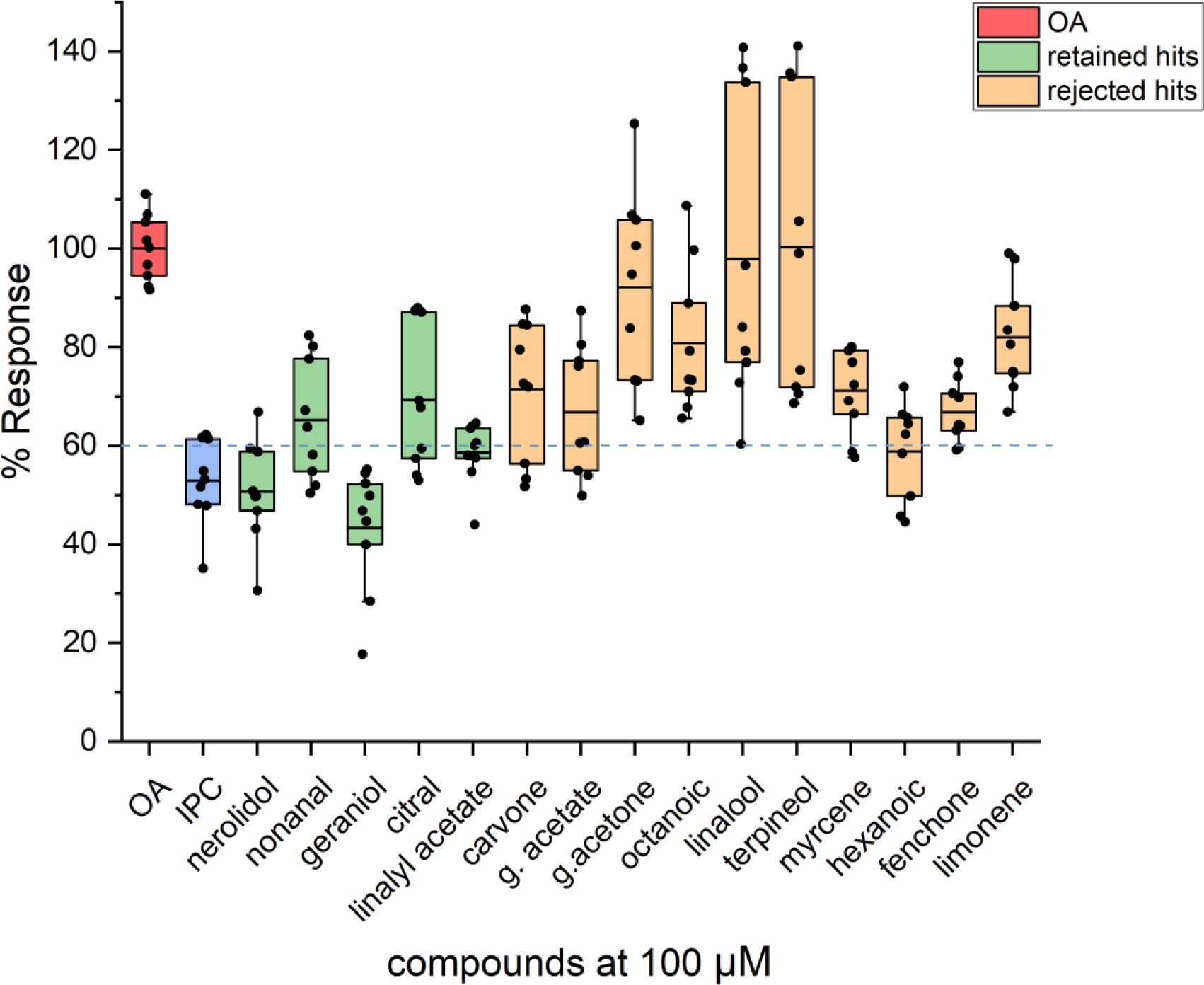
*Ex vivo* functional validations. Assays were carried out for 15 randomly selected compounds that included 5 workflow-retained (green) and 10 workflow-rejected (orange) compounds. The Orco/Photina activity platform (Kythreoti et al., 2021; Tsitoura et al., 2015, 2016) was employed using isopropyl cinnamate (IPC; blue) as antagonist activity standard (52% response or 48% inhibition of the normal activity in the presence of 100μM OA). The primary compound additions, each at a 100μM concentration, did not induce significant ORco channel function (<15% for all of them; data not shown). The cutoff response point for antagonistic activity against 100μM OA was arbitrarily set at <60% (>40% inhibition of the 100% activity obtained by addition of 100μM Orco plus solvent shown in red). Box plots depict mean values, with upper and lower quartiles, and the range of outliers within 1.5IQR are indicated by error bars. The response values for the tested compounds are listed in **Table 6**.

**Table 6.**
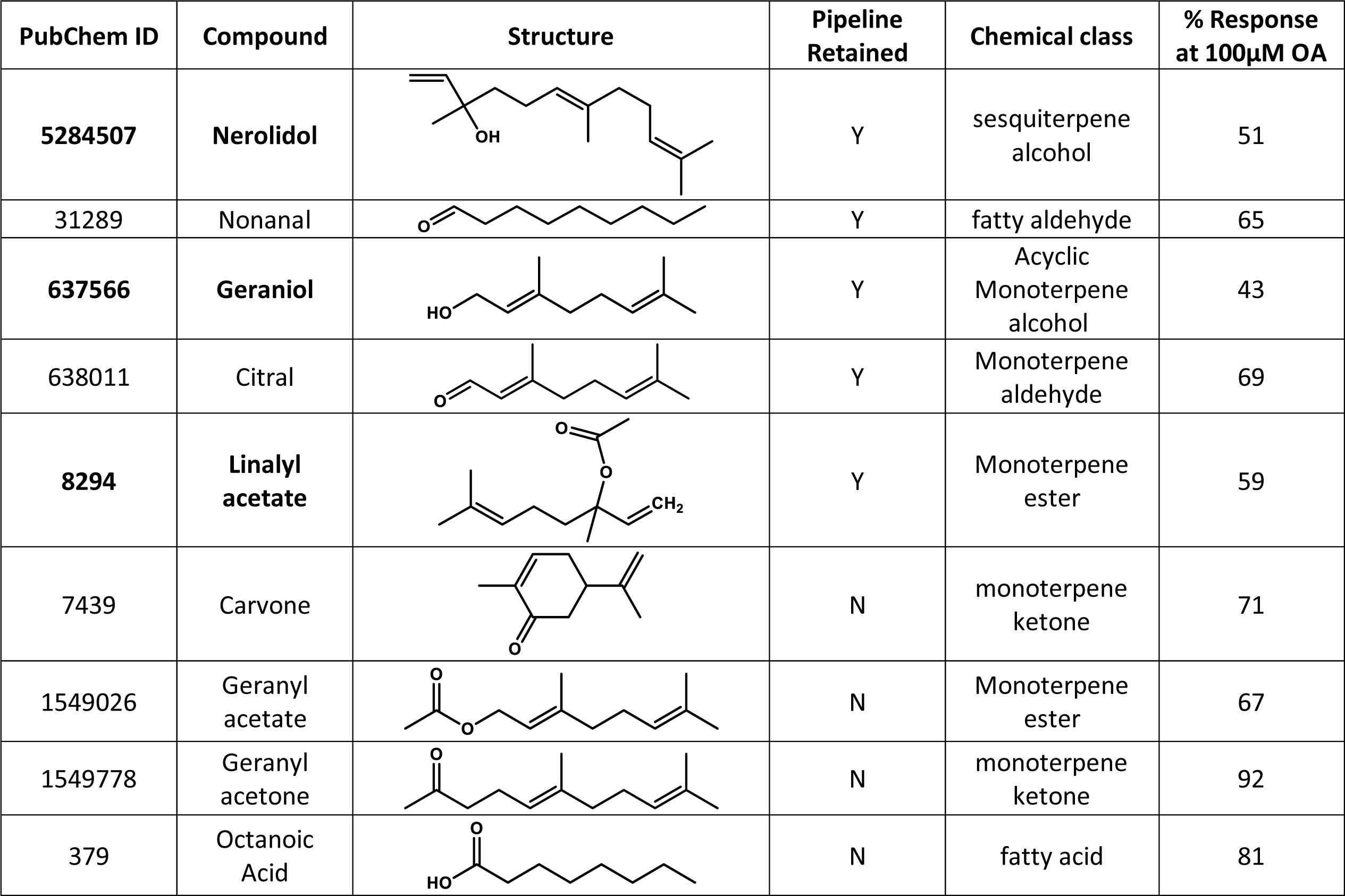
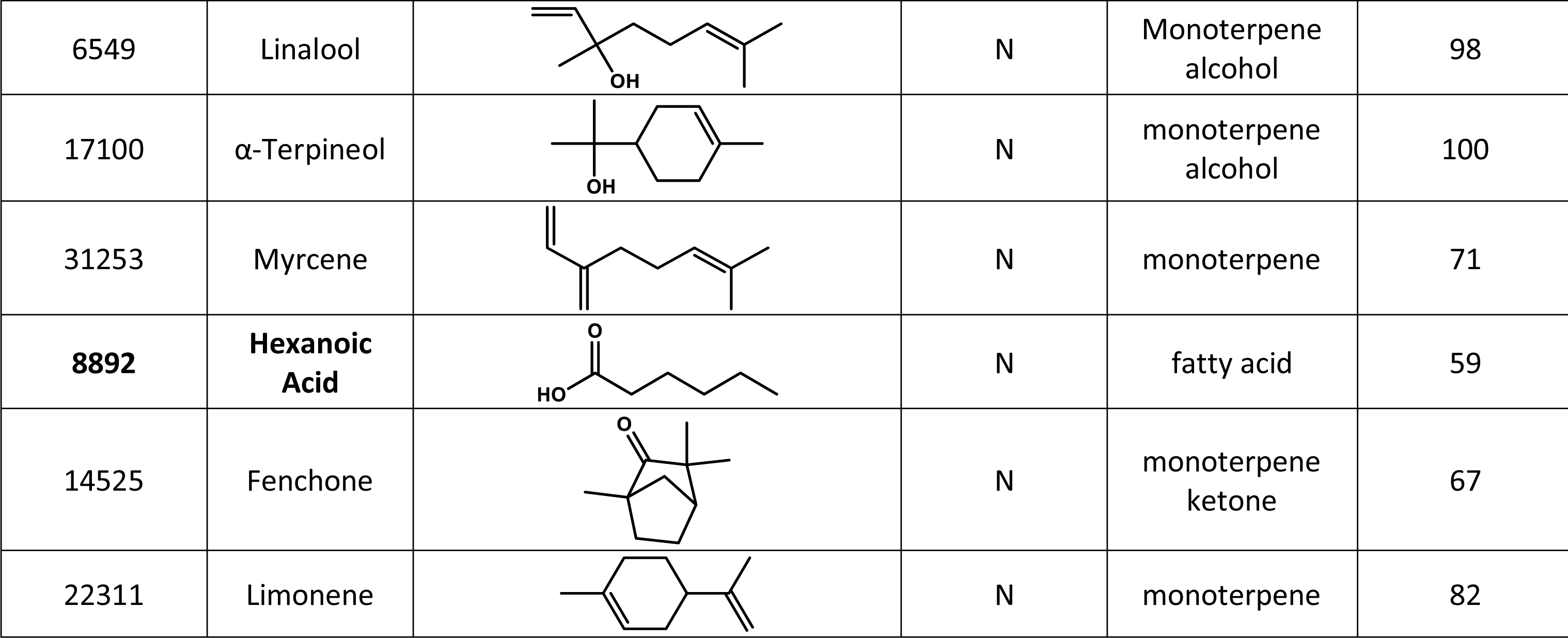
Selected Pharmacophore-SVM pipeline hits and initial functional testing. Compounds causing ≤60% response are indicated in bold.

| PubChem ID | Compound | Structure | Pipeline Retained | Chemical class | % Response at 100μM OA |
| --- | --- | --- | --- | --- | --- |
| **5284507** | **Nerolidol** |  | Y | sesquiterpene alcohol | 51 |
| 31289 | Nonanal |  | Y | fatty aldehyde | 65 |
| **637566** | **Geraniol** |  | Y | Acyclic Monoterpene alcohol | 43 |
| 638011 | Citral |  | Y | Monoterpene aldehyde | 69 |
| **8294** | **Linalyl acetate** |  | Y | Monoterpene ester | 59 |
| 7439 | Carvone |  | N | monoterpene ketone | 71 |
| 1549026 | Geranyl acetate |  | N | Monoterpene ester | 67 |
| 1549778 | Geranyl acetone |  | N | monoterpene ketone | 92 |
| 379 | Octanoic Acid |  | N | fatty acid | 81 |
| 6549 | Linalool |  | N | Monoterpene alcohol | 98 |
| 17100 | α-Terpineol | | N | monoterpene alcohol | 100 |
| 31253 | Myrcene |  | N | monoterpene | 71 |
| **8892** | **Hexanoic Acid** |  | N | fatty acid | 59 |
| 14525 | Fenchone |  | N | monoterpene ketone | 67 |
| 22311 | Limonene |  | N | monoterpene | 82 |

As is shown in Figure 8, 3 out of the 5 retained hits showed >40% inhibitory activities, whereas the remaining 2, nonanal and citral, displayed reduced activities bordering the arbitrary cutoff inhibition limit. On the other hand, 9 out of 10 workflow-excluded compounds exhibited no or low (≤40%) inhibitory activity.

A summary of the overall structural properties and *ex vivo* functionality of the validated compounds selected from the *in silico* screening is shown in **Table 6**.

## DISCUSSION

Vector control has been the principal method of preventing vector-borne infectious diseases for over 100 years and remains highly effective when comprehensively applied and sustained. Many insect species, including mosquitoes, have the potential to transmit a wide variety of pathogens to humans and animals, leading to VBDs with substantial socioeconomic impacts. Given the current climatic changes worldwide, which have brought substantial temperature increases in geographic regions with temperate climates, and the concurrent increases in movements of people due to easier travel conditions, such diseases are spreading at an alarming rate in countries where they were previously absent. Therefore, there is a growing demand for novel, long lasting and environmentally friendly repellents and anosmia-inducing agents. However, the classical research methods for discovery of new protective agents against insect bites, particularly in a spatial context that does not involve direct application on human or animal skin surfaces, is a time consuming and expensive task that prevents the development of novel control measures.

Progress in the rate of discovery of protective agents for humans and domestic animals against various insect disease vectors, particularly mosquitoes, has been achieved relatively recently through the employment of *ex vivo* expression systems developed from cultured amphibian oocytes [35–37], and mammalian [36, 38–42] or insect cell cultures [15, 43–45] upon coupling to relevant bioactivity reporter assays. Further enhancement in the rate of discovery of relevant bioactive compounds has been achieved recently through the exploitation of the seminal discovery that upon *ex vivo* expression, the evolutionarily conserved, obligatory odor receptor co-receptor ORco forms homomeric cation channels [11, 13, 46] whose function may be activated by specific agonists such as VUAA1, OrcoRAM2 and derivatives [11, 13, 47] and inhibited by structurally-related antagonists [10, 12, 13, 48] as well as the demonstrated general inhibition of odorant receptor function *in vivo* as a consequence of specific mutations in the ORco subunit [39, 49–55]. These findings led to the development of the notion that inhibition of the general olfactory functions in insects may also be achieved by the binding of volatile ORco antagonists, preferably of natural origin, to the ORco subunit in nearly all ORx/ORco heteromeric receptor complexes in live insects and produce anosmia-like phenotypes on the targeted species. This notion has been amply demonstrated through the use of some of the *ex vivo* expression-activity detection systems mentioned above in throughput formats and by the concomitant demonstration that the great majority of natural volatiles causing *ex vivo* inhibition of ORco, are capable of inhibiting the olfactory functions in laboratory and field mosquito and sandfly populations in a spatial context [26, 27].

The employment of such throughput systems has allowed a very significant acceleration in the rate of discovery of relevant bioactive compounds by activity screening of small size compound collections, and has also allowed the subsequent classification of the new anosmia-causing antagonists into orthosteric and allosteric classes relevant to the the postulated binding site of the previously characterized non-volatile agonists. However, they are still not adequate by themselves, when the requirements for similar screens of large compound collections are considered in terms of time and material costs. To expedite the search for discovery of relevant bioactive molecules in large compound collections, computational screening methods should be applied as virtual pre-screening tools, which could reduce the number of molecules to be functionally screened *ex vivo* for identification of relevant hits to a reasonable level. *In silico* techniques not only accelerate the discovery of potential compounds, but they also facilitate understanding of relevant biological mechanisms. Toward this goal, in this study we have employed a two-step, ligand-based *in silico* pipeline consisting of a first pharmacophore screening (Step-1) and a subsequent SVM filtering step (Step-2). This pipeline showed highly satisfactory performance in predicting active orthosteric antagonists for ORco, as confirmed by follow-up functional validation.

Pharmacophores are frequently used in virtual screening projects, due to their simplicity and their ability to speed up the *in silico* process [18, 56, 57]. Moreover, since they do not depend on specific functional or structural groups, they can identify chemically divergent molecules. The main difficulty in generating successful pharmacophores is to balance their sensitivity and specificity. Therefore, they are commonly combined with other computational techniques such as Support Vector Machines (SVMs), to improve the accuracy of the results [58]. Support Vector Machines are well established in bioinformatics and chemoinformatics, since they can handle high-dimensional data and small datasets and they can model non-linear decision boundaries. They are also adaptable and versatile. However, SVMs can be computationally expensive for large datasets [59, 60]. For this reason, we have employed in our computational screening process, the SVM filtering after the pharmacophore screening step.

Our virtual screening pipeline achieved 0.75 and 0.82 sensitivity and specificity, respectively, resulting in an overall performance of 0.8 for predicting orthosteric antagonists that caused more than 40% inhibition to ORco (Figure 6). Such a performance is notable because elimination of more than 80% of the number of compounds to be tested translates in commensurate time-, material- and labor-consuming costs for *ex vivo* and *in vivo* tests. Thus, our pipeline can both save resources and accelerate discovery of novel agents. Moreover, to the best of our knowledge, our study is the first one that combines a pharmacophore with a SVM model for identification of AgamORco antagonists and specifically orthosteric ones that are advantageous for future site-specific ORco structure-based screening as compared to blind-docking trials.

Our pharmacophore model (Step-1) resembles the model previously proposed by [61]. That model consisted of a hydrogen-bond acceptor site, two aliphatic and one aromatic hydrophobic site. It was successfully used for virtual screening of an in-house compound database that resulted in four new potential insect repellent candidates. Other studies on insect olfactory ligands [62], employed a Laplacian-corrected Naïve Bayesian machine learning, ligand-based, approach to predict novel volatile *Anopheles gambiae* ORco antagonists. Selected hit compounds were further evaluated for their ability to inhibit electrophysiological responses in adult *Drosophila melanogaster* flies and in behavioral attraction assays against D*. melanogaster* larvae. In contrast to our study, the model was not trained to discriminate between orthosteric and allosteric antagonists. Electroantennography (EAG) recordings of two selected hits, 2-tert-Butyl-6-methylphenol (BMP) and Linalyl formate (LF) suggested an allosteric and non-competitive ORco-dependent mechanism, which was further confirmed by concentration-inhibition analysis of BMP in *Xenopus laevis* oocytes expressing AgamORco. Machine learning techniques such as RandomForest and kNN classifier (IBk; reference?) have also been successfully employed to predict new receptor agonists other than ORco i.e., SlitOR24 and SlitOR25 from *Spodoptera littoralis* [63]. A Supported Vector Machine (SVM) model, (such as Step-2 in our pipeline) has been used for identification of agonists for SlitOR25 [64].

While our approach is characterized by high performance, as with any other prediction method, it could not be 100 percent accurate. For example, hexanoic acid that has been rejected by our workflow at the pharmacophore selection step, showed antagonist activities and is therefore considered as a false negative compound (Figure 7). A meta-analysis of the structure-activity relationship of the hits listed in **Table 6**, has revealed that hexanoic has the smaller length (6 carbon atoms) among the linear hits. In its most extended conformation, the distance between the two centroid hydrophobic features Hyd (carbon atoms) is 6.4 Å and do not conform with the pharmacophore model shown in Figure 1, where the optimum Hyd1-Hyd2 distance has been determined to be 7.2 Å. Given that the initial set (**Table1**) as well as the set used for pharmacophore training (**Tables 2 and S2**) are dominated by longer chain linear compounds (8 to 10 carbon atoms), that can obtain conformations satisfying pharmacophore distances as well as bulky cyclic and aromatic compounds, it is possible that the pharmacophore model is negatively biased toward molecules of smaller length. Such inconsistencies of the model could be eliminated by incorporating more experimental data on short-length agonists. Furthermore, *ex vivo* concentration-inhibition analysis remains to be performed to exclude that hexanoic acid cannot act as an allosteric antagonist i.e., that it is a true negative result (as per terminology of our pipeline for the allosteric *ex vivo* active compounds). On the other hand, two compounds, nonanal and citral that have been retained by our workflow (Figure 7), showed borderline activities in *ex vivo* experiments (Figure 8) and are classified as false positives. These two aldehydes can participate in only one hydrogen bond through their carbonyl group (hydrogen bond acceptor), in contrast with the other three active compounds in the series, which can participate in two hydrogen bonding interactions. In particular, nerolidol and geraniol bear a hydroxyl group that can act as a hydrogen bond acceptor/donor whereas linalyl acetate bears an acetate ester with two oxygen atoms in proximity that can act as hydrogen bond acceptors (**Table 6**).

Concerning the pipeline-rejected hits linalool and α-terpineol, both tertiary alcohols of molecular weights 154.25, with very similar cLogP values of 2.468 and 2.369, respectively, and identical PSA 20.23 (SlogP_VSA1=0), despite the high variability of the *ex vivo* obtained response values they are considered no- or low-activity inhibitors (Figure 8). Moreover, the rejected carvone and fenchone, can participate in only one hydrogen bond, while limonene lacks a functional group for participation in hydrogen bonds (**Table 6**). All three compounds are relatively compact cyclic molecules. The observed activities could therefore, upon further investigation, be a result of allosteric binding. Finally, the rejected geranyl acetone, similar to the retained nonanal and citral, can only participate in one hydrogen bond, while myrcene contains no polar functionalities. Geranyl acetate and Octanoic acid have similar KierA2 and SlogP_VSA1 parameters well outside the decision boundaries for active orthosteric antagonists (Figure 7). Hexanoic acid, which does not conform the pharmacophore model (*vide supra*) but lies inside the decision boundaries, has identical SlogP_VSA1 to geranyl acetate and octanoic acid but different KierA2, due to the different spatial density of atoms in this shorter molecule. Similarly, despite their similar KierA2, the inactive geranyl acetate with SlogP_VSA1=7.74, has both larger SPA (26.3) and more hydrophobic character (cLogP 3.264) than the active geraniol with SlogP_VSA1=0 (SPA=20.23 and cLogP=2.524).

We are noting that compound #88 [ethyl (E/Z)-2-(cyclohex-2-en-1-ylidene) acetate], the single orthosteric antagonist that caused minimal behavioral effects at the dose of 10 nmole/cm^2^, was found to have the highest IC_50_ (195.7 μΜ) among its active counterparts against the activity of 100μM of the ORco agonist OrcoRAM2 in the *ex vivo* tests (**Tables 3** and **5**). Therefore, its low *in vivo* activity may be due to its low inhibitory potency against ORco or/and its relatively high calculated volatility (VP= 0.207 mmHg), that might affect its performance under the 5-min experimental timescale of the behavioral assays. Concerning its weak *ex vivo* binding to ORco and its low *in vivo* activity, it should also be noted that this compound has been tested as a mixture of E/Z isomers. It is very likely that ORco selectively binds one of the two isomers as has been shown be the case for compounds binding to other olfactory receptors [65, 66]. In support of this notion, **Figure S2** showcases the explicit orientations of either the sp^2^ or the sp^3^ hybridization carbons towards the spheroid F4 (Figure 1). Among the two isomers, the Z is better fitting the specific pharmacophore model since the saturated carbons bearing two hydrogens orient to the larger spheroid F2, whilst the unsaturated (sp^2^) carbon with its one hydrogen is oriented to spheroid F4 providing better occupancy. Moreover, the cyclohexene ring carbons holding a sp^3^ hybridization are also bended, thus contributing to the model complementarity in this isomer. Hence, it is possible that the E isomer is a weak or a non-ORco binder, resulting in the apparent weak inhibitory and behavioral activity of the mixture. While the test of the individual isomers is beyond the scope of this study, the ORco specialization against multiple geometric, diastereomeric or enantiomeric isomers of an olfactory ligand is worth investigating in future studies. Such information can reveal the role of ORco on the remarkable selectivity of insect olfaction and be further exploited in ORco-based *in silico* and *ex vivo* screening approaches.

We also note that compound #74 (Bisabolene; also a mix of isomers) that was found to be active in the *ex vivo* screens (Figure 2), escaped detection by the pharmacophore (**Table 3**). Nevertheless, subsequent analyses showed it to have values placing it within the SVM boundaries (KierA2 = 5.4685, SlogP_VSA1 = 0) and also be marginally active in the *in vivo assays* at a dose of 50 nmole/cm^2^ (data not shown). Accordingly, based on the results of the initial pharmacophore screen, we consider it to be a false negative result of our screening pipeline. Moreover, compounds #39 (2,4-Octadienal), #54 [(*Z*)-3-Nonen-1-ol], #74 (Bisabolene) and #77 (α-Bisabolol) and the allosteric antagonist #62 (α-Pinene oxide), which exhibited mild repellent activities (RI 30%-50%) at the same dose (data not shown), were not tested at the lower dose of 10nmole/cm^2^. Future studies should aim to include a more comprehensive evaluation of all *ex vivo*-tested compounds to determine their minimum effective doses and thus provide a more complete understanding of their structure-activity relationships.

To conclude, any ligand-based approach is bound to exhibit some limitations. The shape and electrostatic potential of the ORco binding site and the conformation, hydrophobicity, polarity, and hydrogen bonding potential of the interacting amino acid residues are the determining factors for discrimination of even subtle differences in physicochemical properties and active conformations between inactive compounds and physiologically relevant ligands. Nevertheless, our pipeline was successful in predicting the presence of at least two strong AgamORco orthosteric antagonists, Nerolidol and Geraniol and also confirming the presence of a third one, Linalyl acetate, that had been identified previously as such (Kythreoti et al., 2021), in the collection of 241 odorant compounds, and thus asserting its validity as a screening tool for accelerating discovery of AgamORco orthosteric antagonists. Future, optimization by incorporation of more experimental data could significantly improve its performance. Furthermore, given the availability of 3D-structures of the *A. bakeri* ORco and the structural homologue MhOR5 [8, 9], reliable AgamORco homology models of apo- and liganded form can be created [29] and combined with our *in silico* ligand-based pipeline and *ex vivo* evaluation platform. To this end, our pipeline can constitute the first step for screening large chemical libraries and proposing candidates for subsequent site-specific molecular docking and MD simulations against AgamORco homology models. Such an approach is currently underway for seeking both novel active compounds and gaining structural insights on ligand recognition mechanism by AgamORco.

## MATERIALS & METHODS

### Pharmacophore model development

Based on the previously published orthosteric antagonists and inactive or allosteric compounds, several pharmacophore models were developed using Molecular Operating Environment software (MOE v. 2016.0801; Chemical Computing Group Inc., 1010 Sherbrooke St. West, Suite #910, Montreal, QC, Canada, H3A 2R7, 2016). The Unified annotation scheme was employed including H-bond Donors and Acceptors, as well as Hydrophobic Atoms and Hydrophobic Centroids. The radius of all features was set to 1Å, except for the radius of Hydrophobic Atoms, which was set to 0.7 Å. Query Spacing and Active Coverage were set to 0.9 and 1 correspondingly. Therefore, the generated pharmacophore models were required to match all orthosteric input molecules, while keeping the number of false positives to a minimum. The selected pharmacophore model was used to screen a collection of small molecules of natural origin to identify orthosteric ORco antagonists.

Sensitivity, Specificity and Power Metric (PM) [31] were used for the evaluation of virtual screening performance. They are defined as,

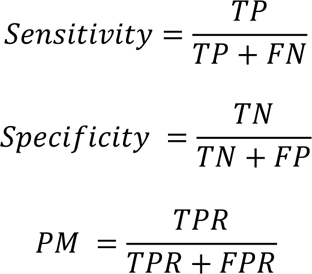

where TPR= TP/(TP+FN) and FPR = FP/(FP+TN) are the true positive rate and the false positive rate correspondingly, TP the number of true positives, FN the number of false negatives, FP the number of false positives and TN the number of true negatives. PM ranges from 0 to +1. Values around 0.5 indicate poor to random models, whereas values between 0.9 and 1.0 are calculated for high quality models. PM is statistically robust with respect to the ratio of actives to the total number of compounds and can be safely applied in early-recognition virtual screening problems.

To improve the performance of the pharmacophore model and to further understand the key features of orthosteric antagonists, we implemented the following procedure. Using MOE, we calculated all the 2D QuaSAR-Descriptors of the molecules identified by the selected pharmacophore model. For each combination of two calculated descriptors, we generated in MATLAB a support vector machine (SVM) with a Gaussian or radial basis function kernel, to classify the orthosteric antagonists from the rest of the molecules. The SVM with the lowest out-of-sample misclassification rate was subsequently optimized and the ten-fold cross-validation loss was reported.

### Chemicals

Compounds analyzed in this study, VOCs and known repellents are presented in **Tables S1 and S2.** Carvacrol (CRV, **I**), linalyl acetate (LA, **4**), (2E,4E)-2,4-octadienal (OCT, **39**) and ethyl cinnamate (EC, **IV**) were purchased from Sigma Aldrich; isopropyl cinnamate (IPC, **II**) from Alfa Aesar; cumin alcohol (CA, **III**) from Acros Organics; N-(4-ethylphenyl)-2-[[4-ethyl-5-(3-pyridinyl)-4H-1,2,4-triazol-3-yl]thio}acetamide (ORco Receptor Agonist ORcoRAM2; OA) from Asinex Corporation and Vitas M Chemical Ltd; *N,N*-diethyl-3-methylbenzamide (DEET; **V**) from Sigma-Aldrich; and coelenterazine from Biosynth. All other VOCs were provided by the Institute of Organic Chemistry, Technische Universität Braunschweig, Germany. For the insect cell-based screening assay, initial stock solutions were prepared as needed and stored at −20 °C. The ORco agonist ORcoRAM2, stocks were prepared in dimethyl sulfoxide (DMSO) whereas the VOCs and coelenterazine stocks were prepared in ethanol. The assay was performed in modified Ringer’s buffer (25 mM NaCl, 190 mM KCl, 3 mM CaCl2, 3 mM MgCl2, 20 mM Hepes, 22.5 mM glucose, pH 6.5; 35), so that the final concentration of DMSO used not to exceed the range of 0.2% to 0.35%.

### Transformation of Bm5 cells for AgamORco and Photina expression and Ca^2+^ influx assays

An insect cell-based assay was employed as a screening platform for the identification and analysis of novel ORco ligands capable of modifying olfaction-mediated mosquito behaviors. Lepidopteran cultured cells (Bombyx mori Bm5; [67], constitutively expressing the AgamORco ligand-gated ion channel were employed, along with a reporter photoprotein Photina [68]. Briefly, Bm5 cells were transformed to stably express cDNAs for AgamORco and Photina from high-expression-level pEIA plasmid vectors as previously described [44, 69–71]. Upon ligand binding activation of the ORco channel, Ca^2+^ ions entering the cells in turn activate the photoprotein, resulting in an increase in luminescence. Cell lines were grown in IPL-41 insect cell culture medium (Genaxxon Bioscience GmbH) supplemented with 10% fetal bovine serum (Biosera) in the presence of 10 μg/ml puromycin and maintained at 28 °C. The ligand binding to the ORco channel and subsequent functional effects were monitored via luminescence emission of the Ca^2+^ influx Photina biosensor, as previously reported [14, 72]. Specifically, insect cells resuspended in modified Ringer’s buffer were seeded in white 96-well plates (200,000–300,000 cells/well), and incubated at room temperature in the dark with 5 μM coelenterazine. Luminescence emissions were then recorded in an Infinite M200 microplate reader (Tecan) at 4s intervals for up to 20s, using buffer and 1% Triton-X100 as baseline and maximum intensities, respectively. Tested compounds were initially added at a 100 μΜ final concentration and the ORco channel response was monitored for 10s at 4s intervals. Cells were allowed to return to baseline, allowing for the monitoring of the secondary effect of ligand binding (4s intervals for 80s), resulting from the addition of 100 μM OA activating the ion channel. Luminescence data were acquired using i-Control 1.3 software by Tecan and normalized by considering ORco agonist luminescent response as the maximal (100%) receptor response for each experimental set. Independent experiments were run in triplicate and repeated at least three times.

### Binding assays

ORco response inhibitions of identified antagonists were further analyzed to determine orthosteric or allosteric binding, relative to the OA (ORcoRAM2) binding site. Solvent or identified antagonists were added to insect cells, constitutively expressing AgamORco and Photina, at concentrations ranging from 1 μM to 1 mM. A 96-well format assay was also employed as described above, and the induced luminescence, if any, was measured. Subsequent addition of OA at different concentrations, 50, 100 or 150 μM were carried out as antagonist dose-dependent inhibition assays, illuminating the type of ligand binding on ORco. OriginPro 8 software, by OriginLab Corporation, was used for curve fitting and IC_50_ value calculations. Dose–response curves were plotted by fitting the normalized data into the equation, where A_1_ and A_2_ are the bottom and top asymptotes, respectively, p is the Hillslope, y is the percent response at a given concentration, and x is logarithm of ligand concentration. Independent experiments were run in triplicate and repeated at least three times.

### Laboratory rearing of Aedes albopictus

Adult *Ae. albopictus* mosquitoes were obtained from the laboratory colony of the Benaki Phytopathological Institute (Kifissia, Greece). The colony is maintained under specific laboratory conditions (25 ± 2 °C, 80% relative humidity, and a 16/8-hour light/dark photoperiod). Larvae were reared in cylindrical enamel pans filled with tap water, with approximately 400 larvae per pan. They were fed ad libitum with powdered fish food (JBL Novo Tom 10% Artemia) until they emerged as adults [26].

### Repellence Bioassays

For the in vivo determination of the repellent activity of tested compounds, the assessment was based on human hand landing counts using cages (33×33×33 cm) equipped with a 32×32 mesh on one side. Each cage contained 100 adult mosquitoes (5 to 10 days old, sex ratio 1:1) starved for 12 hours at 25 ± 2 °C and 70–80% relative humidity [32]. Tested compounds were applied on chromatography paper (Whatman), covering a total area of 24 cm², at dose equivalent to 50 nmole/cm², diluted with dichloromethane (DCM). Data concerning the repellency indices were analyzed using the Kruskal–Wallis test [73]. When significant differences were detected, Mann–Whitney U tests were carried out for pair-wise comparison with a Bonferroni correction for adjustment of p-values [74]. Mosquito landings for each treatment were counted over 5-minute periods. Landing numbers were converted to repellency indices (RI ± SE) using the following equation: RI = [1 -T/C] x 100, where C is the number of landings in the control and T is the number of landings in the treatment [26].

## Supporting information

Supplemental Spreadsheet (Chemical Library of 241 VOCs)

Author contributions statement

## Acknowledgments

The research project was supported by the Hellenic Foundation for Research and Innovation (H.F.R.I.) under the “1^st^ Call for H.F.R.I. Research Projects to support Faculty Members & Researchers and the Procurement of High-and the procurement of high-cost research equipment grant” (Project Number: HFRI-FM17-637).

---

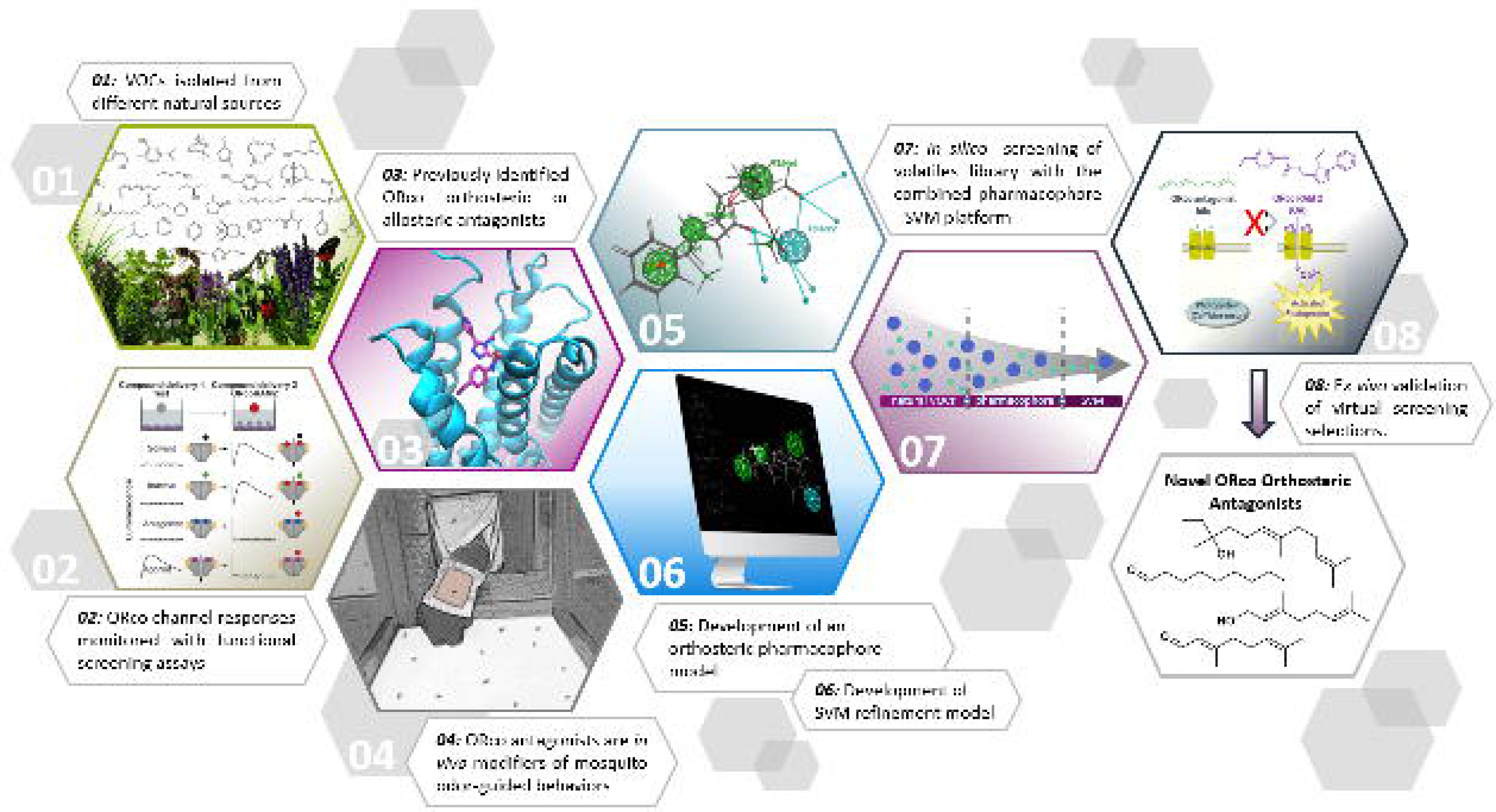

