## Supplemental Information for "Natural volatiles causing inhibition of mosquito biting behaviors: development of a virtual screening platform predicting antagonists of ORco function for accelerated discovery"

^1^Institute of Biosciences and Applications, National Centre for Scientific Research “Demokritos”, Athens, Greece; ^2^Department of Biotechnology, Agricultural University of Athens, Athens, Greece; ^3^Institute of Chemical Biology, National Hellenic Research Foundation, Athens, Greece; ^4^Scientific Directorate of Entomology and Agricultural Zoology, Benaki Phytopathological Institute, 145 61 Kifissia, Greece; ^5^Institute of Organic Chemistry, Technische Universität Braunschweig, Braunschweig, Germany

^b^Department of Science and Mathematics, Deree – The American College of Greece, Athens, Greece.

#Communicating authors:

**Table S1. Training set of 54 VOCs used for development of pharmacophore model.** The set includes 4 orthosteric (green) and 50 negatives; 3 allosteric (orange) and 47 inactive compounds.

| **No** | **Compound** | **Structure** | **CAS No** | **MW** | **Source** |
| --- | --- | --- | --- | --- | --- |
| **I** | **Carvacrol** | 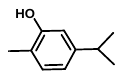 | **499-75-2** | **150.2** | **Plants** |
| **II** | **Isopropyl cinnamate** | 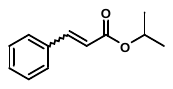 | **7780-06-5** | **190.2** | **Plants** |
| **III** | **Cumin alcohol** | 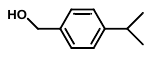 | **536-60-7** | **150.2** | **Plants** |
| **IV** | **Ethyl cinnamate** | 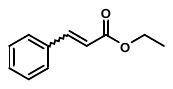 | **103-36-6** | **176.2** | **Plants** |
| **V** | ***N*,*N*-diethyl-3-methylbenzamide (DEET)** | 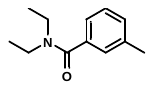 | **134-62-3** | **191.3** | **Synthetic** |
| **1** | 2,5-Dihydrofuran | 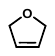 | 1708-29-8 | 70.1 | Plants |
| **2** | 2-Butanone | 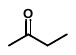 | 78-93-3 | 72.1 | Bacteria |
| **3** | Butyl amine | 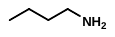 | 109-73-9 | 73.1 | Plants, bacteria |
| **4** | **Linalyl acetate** | 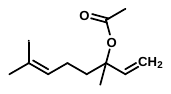 | **115-95-7** | **196.3** | **Plants** |
| **5** | (*S*)-2-Butanol | 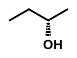 | 4221-99-2 | 74.1 | Bacteria |
| **6** | Pyridine | 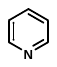 | 110-86-1 | 79.1 | synthetic |
| **7** | Pyrazine | 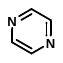 | 290-37-9 | 80.1 | Bacteria |
| **8** | 2-Cyclopenten-1-one | 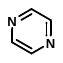 | 930-30-3 | 82.1 | Plants |
| **9** | Thujopsen | 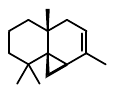 | 470-40-6 | 204.4 | Plants |
| **10** | 3,4-Dihydro-2*H*-pyran | 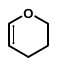 | 110-87-2 | 84.1 | Synthetic |
| **11** | Cyclopropyl methyl ketone | 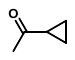 | 765-43-5 | 84.1 | Synthetic |
| **12** | 2-Pyrrolidone | 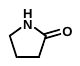 | 616-45-5 | 85.1 | Spiders |
| **13** | 2-Pentanone | 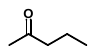 | 107-87-9 | 86.1 | Plants |
| **14** | *N*-Methyl propionamide | 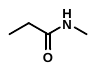 | 1187-58-2 | 87.1 | Algae |
| **15** | 1,3-Butanediol | 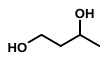 | 107-88-0 | 90.1 | Bacteria |
| **16** | Phenol | 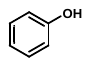 | 108-95-2 | 94.1 | Bacteria |
| **17** | (2*E*,4*E*)-Hexadienal | 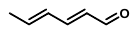 | 142-83-6 | 96.1 | Insects |
| **18** | 3-Methyl-2-cyclopenten-1-one | 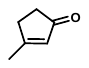 | 2758-18-1 | 96.1 | Bacteria |
| **19** | Furfuryl alcohol | 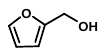 | 98-00-0 | 98.1 | Bacteria |
| **20** | *(E*)-3-Hexen-1-ol | 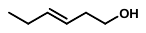 | 928-97-2 | 100.2 | Plants |
| **21** | δ-Valerolactone | 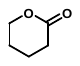 | 542-28-9 | 100.1 | Plants |
| **22** | γ-Valerolactone | 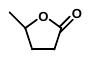 | 108-29-2 | 100.1 | Bacteria |
| **23** | 2-Methyl-4-butanolide | 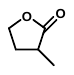 | 1679-47-6 | 100.1 | Fungi |
| **24** | 2,3-Pentandione | 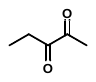 | 600-14-6 | 100.1 | Yeast |
| **25** | Methyl isobutyrate | 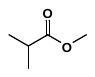 | 547-63-7 | 102.1 | Bacteria |
| **26** | α-Humulene | 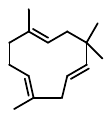 | 6753-98-6 | 204.4 | Plants |
| **27** | 3-Methylthio-1-propanol | 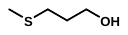 | 505-10-2 | 106.2 | Bacteria |
| **28** | Anisole | 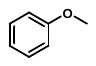 | 100-66-3 | 108.1 | Plants |
| **29** | Benzyl alcohol | 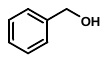 | 100-51-6 | 108.1 | Plants |
| **30** | 2-Acetyl-1H-pyrrole | 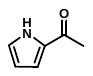 | 1072-83-9 | 109.1 | Plants |
| **31** | 2-Ethylthiophene | 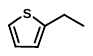 | 872-55-9 | 112.2 | Bacteria |
| **32** | 1,8-Cineole | 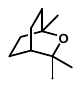 | 470-82-6 | 154.3 | Plants |
| **33** | 2-Heptanone | 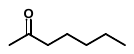 | 110-43-0 | 114.2 | Plants |
| **34** | (*R*)-Sabinene | 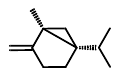 | 2009-00-9 | 136.2 | Plants |
| **35** | Benzyl cyanide | 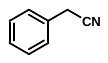 | 140-29-4 | 117.2 | Plants |
| **36** | Phenethyl amine | 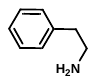 | 64-04-0 | 121.2 | Bacteria |
| **37** | 2-Phenylethanol | 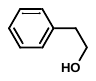 | 60-12-8 | 122.2 | Bacteria |
| **38** | p-Hydroxybenzaldehyde | 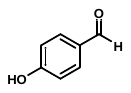 | 123-08-0 | 122.1 | Bacteria |
| **39** | **2,4-Octadienal** | 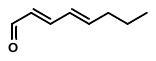 | **5577-44-6** | **124.2** | **Insects** |
| **40** | 6-Methyl-5-hepten-2-one | 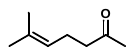 | 110-93-0 | 126.2 | Insects |
| **41** | 2-Acetyl thiazole | 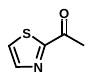 | 24295-03-2 | 127.2 | Bacteria |
| **42** | 4-Octanone | 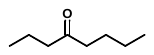 | 589-63-9 | 128.2 | Plants |
| **43** | 2-Octanone | 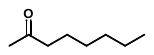 | 111-13-7 | 128.2 | Plants |
| **44** | Ethyl isovalerate | 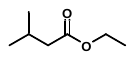 | 108-64-5 | 130.2 | Plants |
| **45** | **(1*S*)-3-Carene (CAR)** | 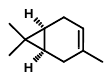 | **498-15-7** | **136.2** | **Plants** |
| **46** | (*R*)-Limonene | 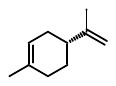 | 5989-27-5 | 136.2 | Plants |
| **47** | (*S*)-Limonene | 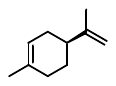 | 5989-54-8 | 136.2 | Plants |
| **48** | Camphene | 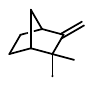 | 79-92-5 | 136.2 | Plants |
| **49** | α-Pinene | 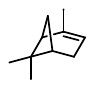 | 7785-26-4 | 136.2 | Plants |
| **50** | β-Pinene | 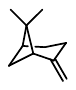 | 18172-67-3 | 136.2 | Plants |

**Table S2: Collection of 49 natural VOCs.** “unseen” set of data, used for pharmacophore screening. Green color indicates pharmacophre hit compounds.

| **No** | **Compound** | **Structure** | **CAS No** | **MW** | **Source** |
| --- | --- | --- | --- | --- | --- |
| **51** | 2′-Hydroxy acetophenone | 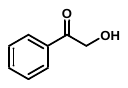 | 582-24-1 | 136.15 | bacteria |
| **52** | Methyl nicotinate | 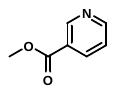 | 93-60-7 | 137.14 | bacteria |
| **53** | 2-Nonanone | 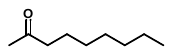 | 821-55-6 | 142.24 | plants |
| **54** | **(*Z*)-3-Nonen-1-ol** | 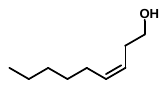 | **10340-23-5** | **142.24** | **Plants** |
| **55** | (*S*)-Perillaldehyde | 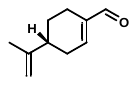 | 18031-40-8 | 150.22 | plants |
| **56** | Methyl anthranilate | 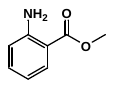 | 134-20-3 | 151.17 | plants |
| **57** | Pulegone | 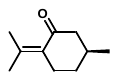 | 89-82-7 | 152.24 | plants |
| **58** | Vanillin | 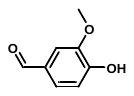 | 121-33-5 | 152.15 | plants |
| **59** | Limonene oxide (cis/trans mix) | 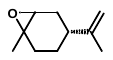 | 203719-54-4 | 152.24 | plants |
| **60** | **(2*E*,4*E*)-Decadienal (DEC)** | 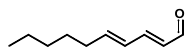 | **25152-84-5** | **152.24** | **Plants** |
| **61** | (1*R*)-(−)-Fenchone | 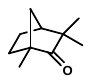 | 7787-20-4 | 152.23 | plants |
| **62** | **α-Pinene oxide** | 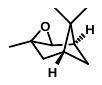 | **1686-14-2** | **152.23** | **Plants** |
| **63** | 2,6,6-Trimethyl-2-cyclohexene-1,4-dione | 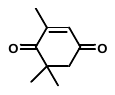 | 1125-21-9 | 152.19 | plants |
| **64** | **Borneol** | 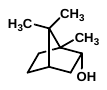 | **464-45-9** | **154.25** | **Plants** |
| **65** | p-Menth-1-en-9-ol | 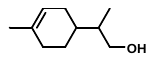 | 18479-68-0 | 154.25 | plants |
| **66** | **2-Methylquinoline** | 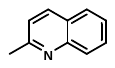 | **612-58-8** | **143.19** | **Bacteria** |
| **67** | *N*-2-Phenyl ethylformamide | 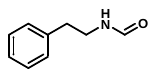 | 23069-99-0 | 149.19 | bacteria |
| **68** | *cis*-Jasmone | 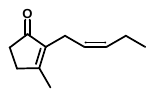 | 488-10-8 | 164.24 | plants |
| **69** | 4-Methyl-1-pentanol | 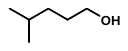 | 626-89-1 | 102.17 | bacteria |
| **70** | Syringaldehyde | 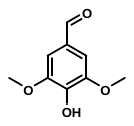 | 134-96-3 | 182.17 | butterflies |
| **71** | γ-Undecalactone | 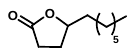 | 104-67-6 | 184.28 | plants |
| **72** | 2-Tridecanone | 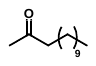 | 593-08-8 | 198.35 | plants |
| **73** | Longifolene | 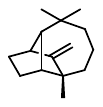 | 475-20-7 | 204.36 | plants |
| **74** | **Bisabolene (mix of isomers)** | 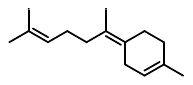 | **495-62-5** | **204.36** | **Plants** |
| **75** | **Aromadendrene** | 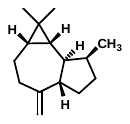 | **489-39-4** | **204.35** | **Plants** |
| **76** | β-Caryophyllen | 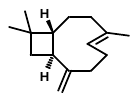 | 87-44-5 | 204.36 | plants |
| **77** | **α-Bisabolol** | 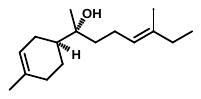 | **515-69-5** | **222.37** | **Plants** |
| **78** | 1-Hexadecanol | 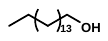 | 36653-82-4 | 242.45 | plants |
| **79** | Phytol | 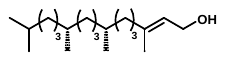 | 150-86-7 | 296.54 | plants |
| **80** | Butyl 2-amino-4-chlorobenzoate | 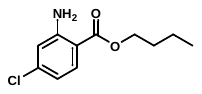 | 173364-37-9 | 227.69 | bacteria |
| **81** | (*Z*)-Octadec-11-enenitrile | 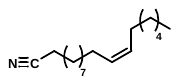 | 2093387-42-7 | 263.47 | bacteria |
| **82** | (*R*)-4-hydroxy-2,6,6-trimethyl-cyclohex-2-en-1-one | 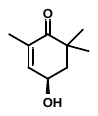 | 76686-09-4 | 154.21 | butterflies |
| **83** | **13-Methyltetradec-3-ene nitrile**  **(cis/trans mix))** | 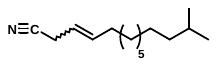 | **2093392-85-7 (Z-isomer)**  **2097108-10-4 (E-isomer)** | **221,39** | **Bacteria** |
| **84** | (9*Z*,12*Z*,15*S*)-Octadeca-9,12-dien-15-olide | 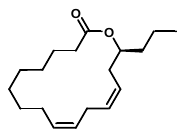 | Not registered | 278.44 | insects |
| **85** | *N*-(3-Methylbutyryl)-*O*-(2-methylpropionyl)-L-serine methyl ester | 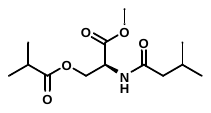 | 2049918-70-7 | 273.33 | spiders |
| **86** | 2-Pentylpyridine | 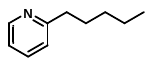 | 2294-76-0 | 149.23 | bacteria |
| **87** | Methyl cyclohex-2-ene-1-carboxylate  (undefined stereochemistry) | 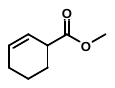 | 25662-37-7 | 154.21 | bacteria |
| **88** | **Ethyl (*E*/*Z*)-2-(cyclohex-2-en-1-ylidene) acetate (cis/trans mix)** | 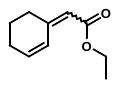 | **136707-84-1**  **71055-14-6**  **(*Z*-isomer)**  **71055-13-5**  **(*E*-isomer)** | **166.24** | **Bacteria** |
| **89** | 7-Tetradecynoic acid | 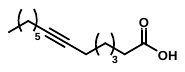 | 55182-86-0 | 224.34 | artificial |
| **90** | 1-Phenylbutane-2,3-dione | 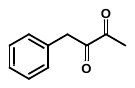 | 38087-02-4 | 162.19 | bacteria |
| **91** | 1,10-Dimethyl-1(9)-octal-2-one  (undefined stereochemistry) | 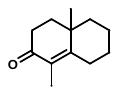 | 54832-12-1  104873-45-2  [[(-)-1](javascript:))]  35493-01-7  [(R)-(+)-1] | 178.28 | bacteria |
| **92** | *N*-Furfur-2-ylisobutylamide | 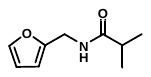 | 24734-02-9 | 167.21 | bacteria |
| **93** | *N*-(2-Phenylethyl) isobutyramide | 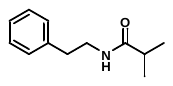 | 71022-62-3 | 191.27 | bacteria |
| **94** | (*R*)-2-Heptyl acetate | 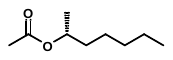 | 54638-12-9 | 158.24 | plants |
| **95** | 2-Pentyl 2-methylbutanoate | 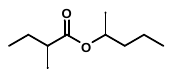 | 57966-40-2 | 172.27 | plants |
| **96** | 13-Methyltetradecane-1-ol |  | 20194-47-2 | 228.42 | bacteria |
| **97** | Linalool oxide (mix of isomers) |  | 60047-17-8 | 168.24 | plants |
| **98** | **(*E*)-3-Methyl-2-(3-methylbutyliden)-4-butanolide** |  | **Not registered** | **198.35** | **Mites** |
| **99** | **(4*R*,6*R*,8*R*)- trimethyldecan-2-one** |  | **158648-77-2** | **136.15** | **Mites** |

**Table S3. Additional data on graphs presented in Figure 3.**

| **Compound** | **OA conc. (μM)** | **IC_50_ (μM)** | **pIC_50_ ± error** | **R^2^** |
| --- | --- | --- | --- | --- |
| **54** | 50 | 26.4 | 4.57856 ± 0.12475 | 0.99999 |
|  | 100 | 48.9 | 4.31096 ± 0.0675 | 0.99999 |
|  | 150 | 76.2 | 4.11814 ± 0.19885 | 0.99997 |
| **60** | 50 | 18.3 | 4.7384 ± 0.05967 | 1 |
|  | 100 | 66.7 | 4.17811 ± 0.38036 | 0.99989 |
|  | 150 | 84.4 | 4.07346 ± 0.14054 | 0.99986 |
| **62** | 50 | 49 | 4.30862 ± 0.25906 | 0.99969 |
|  | 100 | 52 | 4.2837 ± 0.19604 | 0.99969 |
|  | 150 | 64.4 | 4.19083 ± 0.03993 | 0.99998 |
| **64** | 50 | 87.1 | 4.0602 ± 0.24007 | 0.99998 |
|  | 100 | 86.5 | 4.06288 ± 0.09513 | 0.99999 |
|  | 150 | 85.5 | 4.06804 ± 0.09235 | 0.99997 |
| **66** | 50 | 126 | 3.89984 ± 0.14175 | 0.99999 |
|  | 100 | 115 | 3.95015 ± 2.86483 | 0.99996 |
|  | 150 | 110.5 | 3.95157 ± 0.86147 | 0.99998 |
| **75** | 50 | 25.2 | 4.59898 ± 0.09235 | 0.99995 |
|  | 100 | 26.2 | 4.58184 ± 0.18369 | 0.99988 |
|  | 150 | 21.5 | 4.66819 ± 0.07359 | 0.99999 |
| **88** | 50 | 123.9 | 3.90138 ± 0.3215 | 0.99987 |
|  | 100 | 195.7 | 3.71072 ± 0.26149 | 0.99999 |
|  | 150 | 252 | 3.59897 ± 0.25369 | 0.99994 |
| **98** | 50 | 24.2 | 4.61617 ± 0.16337 | 0.99994 |
|  | 100 | 43.2 | 4.36479 ± 0.32587 | 0.99960 |
|  | 150 | 65.4 | 4.18457 | 0.99987 |

**Table S4. Repellency indices of orthosteric and allosteric antagonists.** Data presented in Figure 5 graphs.

| **Compound** | **RI**  **(at 10nmole/cm^2^)** |
| --- | --- |
| **DEET** | 0.84 ± 0.01 |
| **I** | 0.17 ± 0.01 |
| **II** | 0.29 ± 0.04 |
| **60** | 0.71 ± 0.05 |
| **64** | 0.58 ± 0.02 |
| **66** | 0.93 ± 0.01 |
| **75** | 0.25 ± 0.03 |
| **83** | 0.49 ± 0.03 |
| **88** | 0.08 ± 0.02 |
| **98** | 0.55 ± 0.03 |

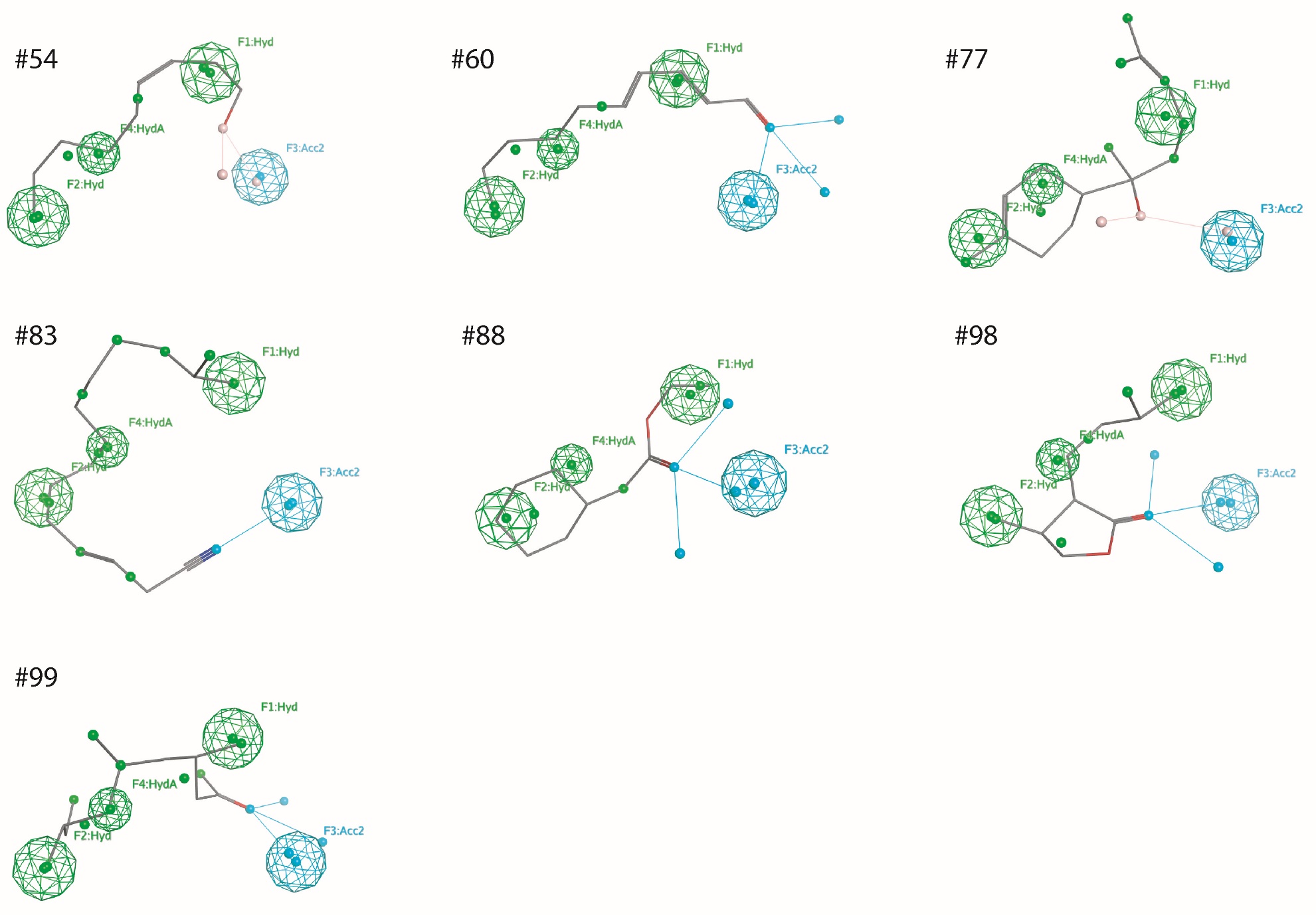

**Figure S1.** Superimposition of *ex vivo* active pharpacophore hits onto the pahrmacophore model

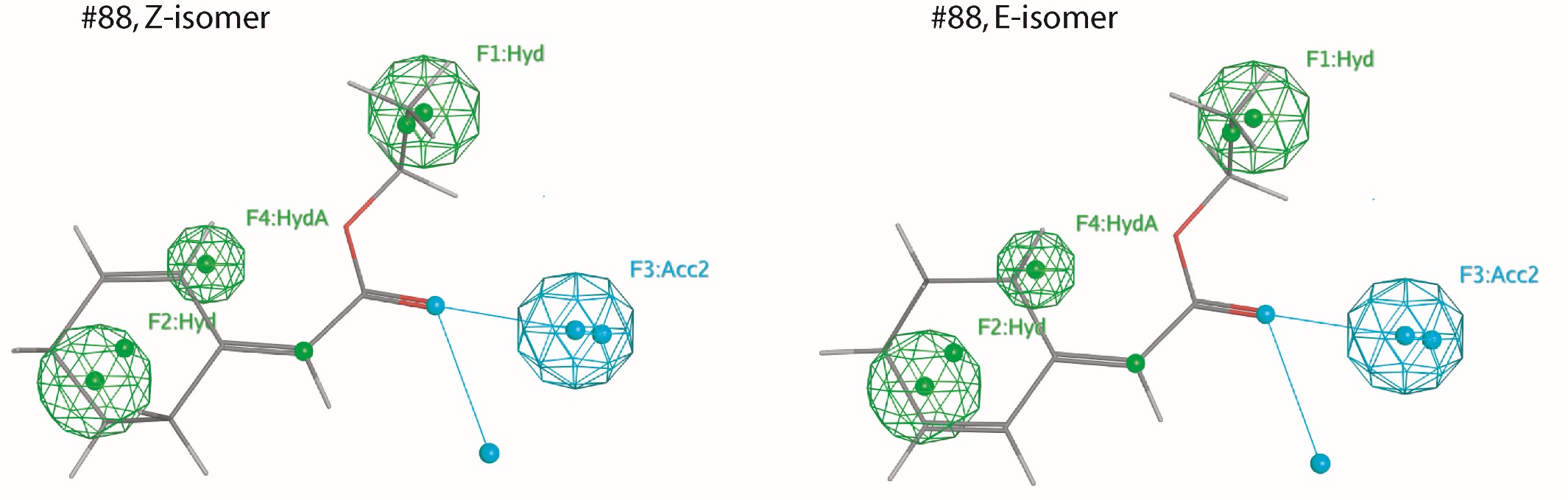

**Figure S2.** Superimposition of *Z* and *E* geometrical isomers of pharmacophore hit #88 onto the pharmacophore model.
