## Supplementary material for "Natural volatiles causing inhibition of mosquito biting behaviors: development of a virtual screening platform predicting antagonists of ORco function for accelerated discovery": Author contributions statement

**Georgia Kythreoti:** Conceptualization of ORco antagonists classes**,** Acquisition, analysis and interpretation of *ex vivo* binding data, **Trias Thireou:** Conceptualization realization and interpretation of *in silico* experiments, **Christos Karoussiotis:** ORco Molecular genetics experimentation**, Zafiroula Georgoussi:** Provided input and critical feedback for ORco genetics and helped shape the research, **Panagiota GV Liggri** and **Vasileios Karras:** Mosquito rearing and Realization of *in vivo* repellence bioassays, **Dimitrios P Papachristos** and **Antonios Michaelakis:** Design of *in vivo* repellence bioassays, Statistical analysis and interpretation of *in vivo* repellence bioassays data, **Stefan Schulz:** Provided compound collections, helped shape the research and gave critical feedback, **Spyros E Zographos:** Provided critical feedback and helped shape the research, Interpretation of *in silico* experiments, Structural analysis and funding acquisition, **Kostas Iatrou:** Conceived and planned the experiments, interpretation of the results and funding acquisition. **All the authors:** writing, review and editing of the manuscript. All authors have read and agreed to the published version of the manuscript.
